# VEGF-A/C co-stimulation, without shear stress, triggers the polarization of lymphatic microvessels

**DOI:** 10.1101/2025.08.13.670220

**Authors:** Jules Edwards, Florent Morfoisse, Baptiste Alric, Barbara Garmy-Susini, Yukiko T. Matsunaga, Aurélien Bancaud

## Abstract

The lymphatic system maintains interstitial fluid homeostasis and supports immune function through dynamic regulation of its architecture mediated by molecular signals—such as vascular endothelial growth factors (VEGFs)—and physical cues. While VEGFs are known to promote endothelial proliferation, their broader roles in tissue organization remain under investigation. Using a lymphatic vessel-on-a-chip platform, we examine how lymphatic endothelial cells (LECs) respond to VEGF-A, VEGF-C, or their combination. We find that co-stimulation synergistically enhances lymphangiogenic sprouting while preserving barrier integrity. Co-stimulation also induces axial polarization of the tissue along the vessel axis, even in the absence of external mechanical stimuli. This polarization requires activation of the VEGFR2/VEGFR3 heterodimer and is disrupted by inhibition of the Src-dependent mechanotransduction pathway. Further co-stimulation enhances LEC motility and triggers vessel contraction. Modeling suggests that the tubular geometry of the lymphatic monolayer imposes intrinsic mechanical anisotropy—softer in the circumferential than axial direction. This geometrically-encoded stiffness landscape directs cell migration along the stiffer axis, uncovering a form of durotaxis driven by curvature-induced anisotropy. These results highlight a previously unrecognized mechanism by which biochemical and biophysical cues direct lymphatic tissue polarization, offering new insight into how geometry and mechanosensing shape lymphatic function.

## INTRODUCTION

The lymphatic system is a unidirectional vascular network, which drains excess interstitial fluid from tissues and returns this liquid, referred to as the lymph, to the blood circulation through the subclavian vein ^1^. Disregarding tissue-specificities of the lymphatic endothelium ^2^, two anatomically distinct lymphatic vessel (LV) structures have been historically described based on their function in lymph drainage. The initial LVs collect the interstitial fluid through loose and discontinuous cell-cell “button junctions” ^3^. Their typical diameter spans 20 to 70 µm, and they are organized in branched architectures ^4^. The lymph is then drained through the collecting LVs, which are characterized by larger diameters of 100 to 500 µm and more impermeable “zipper junctions” with straight intercellular contacts ^5^. The collecting lymphatic vessels also contain intraluminal valves, which prevent lymph backflow as the fluid progresses through the network of collecting vessels and lymph nodes ^6^.

This multiscale architecture results from temporally-concerted reorganizations of the vascular system during development. In mice, the lymphatic vasculature is initially formed at approximately E9.5, when a population of endothelial cells become lymphatic endothelial progenitor cells ^7^. Stimulation by the vascular endothelial growth factor C (VEGF-C) then induces lymphatic vessel sprouting ^8^, in a process termed lymphangiogenesis. The specialization of initial and collecting lymphatic occurs at E16.5-17.5 ^9^, roughly concomitantly with the formation of lymphatic valves ^10^. This latter process involves the dynamic clustering of lymphatic endothelial cells (LECs) to form a ring-like ordered monolayer around the vessel ^10–12^. This morphological change has been thoroughly investigated, showing that its activation involves the integration of biochemical and mechanobiological signaling. Indeed, VEGF-C signaling plays a pivotal role in initiating lymphatic morphological reorganization ^13^, which in turn establishes a feedback loop with the lymphatic-specific transcription factor PROX1 ^14^. In addition, fluid shear stress and mechano-transduction pathways have been shown to contribute to lymphatic tissue reorganization during valve formation ^11,12^. These claims have been supported by cell polarization in 2D culture systems exposed to fluid shear stress. These experiments showed different modes of polarization characterized by parallel or perpendicular orientation relative to the flow direction ^10,15–17^. Furthermore, it has been suggested that LECs’ sensitivity to shear stress is coupled to the level of VEGFR3 expression ^18^, the main VEGF-C receptor ^19^. Whether VEGF-C stimulation alone contribute to LEC polarization within a vessel-like organization remains, to the best of our knowledge, poorly described. This remains unknown because most *in vitro* experiments are performed on flat substrates, *i.e*., overlooking the geometrical specificities of lymphatic vessels, that can for instance induce large-scale ordering of endothelial tissues ^20^.

In this report, we use lymphatic microvessel platforms (LµV) ^21–23^, which are tubular, three-dimensional monolayers of LECs, to assess the effects of two key vascular endothelial growth factors, VEGF-C and VEGF-A, delivered individually or in combination. This experimental design is intended to replicate physiological conditions in which VEGF-A and VEGF-C are often co-expressed, and LECs are typically exposed to multiple VEGF family members during both embryonic development and in adult tissues ^24,25^, meanwhile their potential or synergistic effects are poorly studied. VEGF-A is included not only for its central role for blood vessel angiogenesis via VEGFR-2 interaction ^26^, but also for its lesser, yet documented, contribution to lymphangiogenesis ^24^. By comparing the structure and function of LµVs under control conditions, and following VEGF-A, VEGF-C, or their combination, we first report that co-stimulation synergistically enhances lymphatic sprouting. We then demonstrate that VEGF-A-induced destabilization of the lymphatic barrier is mitigated by co-treatment with VEGF-C. Further VEGF-A/C co-stimulation induces polarization of the lymphatic tissue along the longitudinal axis of the tube. This polarization is dependent on VEGFR-2/VEGFR-3 heterodimerization, as it is abolished by a VEGF-C mutant that selectively targets VEGFR-3 homodimers but not heterodimers. Inhibitor studies confirm the role of VEGFR-2/VEGFR-3 signaling, because activation of the AKT/PI3K mechanotransduction pathway appears to be required for polarization, together with the ROCK kinase, which regulates actin stress fiber formation. Finally, we investigate the biophysical mechanisms underlying this tissue polarization. We find that polarization correlates with increased LEC motility and contractile force generation. Finite element simulations reveal that the tubular geometry of the LµV produces stiffness anisotropy between the axial and circumferential directions, guiding LEC migration along the stiffer axial direction of the vessel. This behavior is consistent with a mechanism of anisotropic durotaxis, driven not by spatial stiffness gradients but by geometric anisotropy. We conclude by discussing how these findings provide insight into the dynamic regulation of lymphatic function and remodeling.

## RESULTS AND DISCUSSION

### VEGF-A/C co-stimulation enhances lymphatic sprouting

We used a tissue engineering approach to fabricate a tubular scaffold in collagen gels of 200 µm in diameter and 6 mm in length ^27–29^ (Fig. 1A; detailed fabrication methods in the Methods section). Using Human Dermal LECs, which represent a standard for *in vitro* studies of lymphatic barrier function ^16,23,30^ and lymphangiogenesis ^21,31,32^, we seeded the cylindrical scaffolds to produce a LµV, that could be cultured for several days. The resulting LEC monolayer expressed the lymphatic vessel endothelial hyaluronan receptor 1 (LYVE-1, Suppl Fig. S1) and vascular endothelial cadherin (VE-Cad; see below).

We focused on the response of LµVs to VEGF-A, VEGF-C, and their combination, each supplemented in the culture medium at a concentration of 100 ng/mL. The choice of concentration aligns with previous microfluidic studies of lymphangiogenesis, where sprouting typically becomes apparent after four days of stimulation ^31,33,34^. To capture early morphological changes, we monitored vessel structure over the first three days using brightfield microscopy. Within this time window, VEGF-A or VEGF-C alone induced little to no sprouting, whereas co-stimulation led to a pronounced onset of sprouting (Fig. 1B). To quantify this effect, we extracted the contour of the LµV (see Methods) and measured its change after 66 hours normalized to the initial contour registered 3 hours post fabrication (Fig. 1C). The relative contour length increased by 24 ± 9% in response to VEGF-A/C, showing a statistically significant difference compared to the control and single growth factor treatments with VEGF-A (10 ± 7%), VEGF-C (4 ± 3%), and control (7 ± 8%; Fig. 1D). Notably, unlike previous reports ^34^, we observed minimal sprout formation under VEGF-C stimulation due to the short time and absence of spatial concentration gradient in our experimental setup. Further, we monitored the formation of sprouts during the first 12 hours of the co-stimulation, and observed a dynamic process with protrusions that either remained stable in the collagen matrix or disappeared due retraction events (Fig. 1E). After 3 days, the architecture of the sprouts, as inferred from immunoconfocal microscopy of DNA and actin, suggested a tip/stalk cell organization with a single tip cell at the leading edge of the sprout and stalk cells organized behind the tip cell (Fig. 1F). Therefore, our data show the accelerated formation of lymphatic sprouts with the synergetic co-stimulation with VEGF-A/C. Moreover, the cellular density within the LµV, estimated by quantifying nuclei in immunoconfocal micrographs (see representative images below), reflected the anticipated proliferative response to endothelial growth factors ^35^.

Specifically, VEGF-A induced a moderate increase in cell density, which was further elevated with VEGF-C stimulation. Co-stimulation with both VEGF-A and VEGF-C further enhanced LEC growth. The measured cell densities were 2.1 ± 1.2, 2.6 ± 1.7, and 3.2 ± 0.7 × 10⁻³ cells/µm² for VEGF-A, VEGF-C, and VEGF-A/C co-treatment, respectively, compared to 1.5 ± 0.3 × 10⁻³ cells/µm² under control conditions (Suppl. Fig. S2). Altogether, stimulation of LECs by endothelial growth factors within LµVs recapitulate the expected enhancement of both lymphangiogenic sprouting and cellular proliferation.

**Figure 1:**
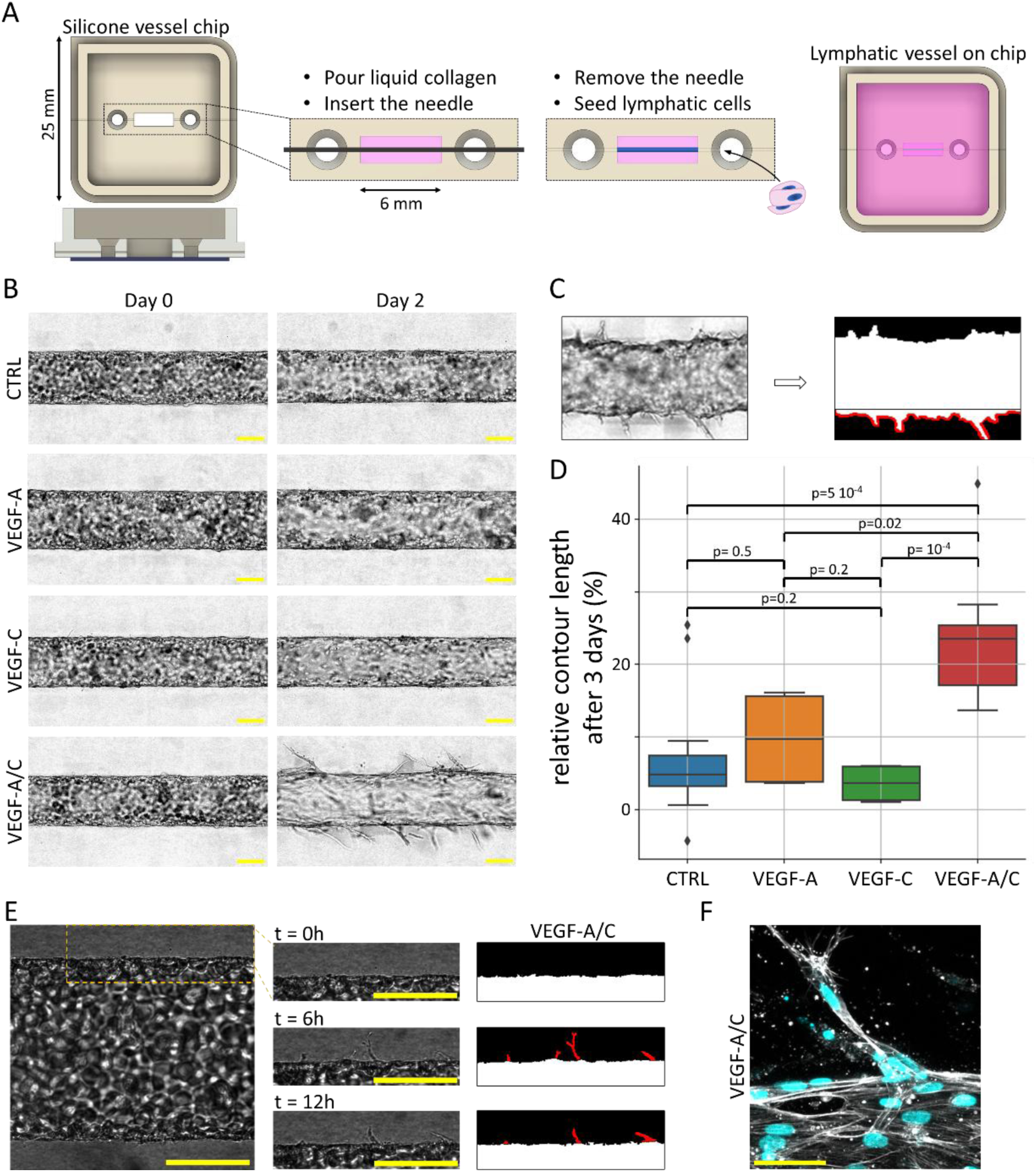
Fabrication of LµV and structural characterization in response to growth factor stimulation. **(A)** Scheme of the silicone LµV shown in grey, and the fabrication protocol to produce a hollow tube in collagen gels represented in pink. After LEC seeding, the endothelial tissue is cultured for three days. **(B)** Optical micrograph of the LµV just after its fabrication and after two days of culture with or without stimulation with growth factors. **(C)** The LµV contour shown in red is segmented on each side of the vessel. **(D)** The graph shows the relative change of the contour length after 3 days of culture with or without growth factor stimulation. **(E)** Live imaging of a LµV shows the dynamic formation of sprouts in response to VEGF-A/C co-stimulation. Segmented sprouts are shown in red. **(F)** Maximum intensity projection (MIP) of confocal micrographs showing a sprout after 3 days of VEGF-A/C co-stimulation. Labels: phalloidin in gray and nucleus in cyan. Scale bars indicate 100 µm, except for the MIP, where it represents 50 µm.

### VEGF-A/C co-stimulation reverse VEGF-A-induced destabilization of the lymphatic barrier function

Lymphatic endothelial barriers constitute barriers to and regulators of solute transport, which are known to be challenged by vascular growth factors ^36^. We thus investigated the LµV barrier integrity in the four different conditions, and correlated this data with the structure of intercellular junctions, as inferred from the VE-Cad immunostaining pattern. The barrier function was quantified using diffusive permeability, which measures how easily molecules passively diffuse across the voids in a cell monolayer ^37^. We used 70 kDa fluorescent dextran (∼12 nm diameter) as a tracer ^38,39^. This dye was injected into the LµV lumen following the installation of a plastic device that maintained constant hydrostatic pressure in the lumen and in the surrounding collagen gel (black solid line in Fig. 2A). Fluorescence redistribution around the vessel was recorded for 5 minutes at 15-second intervals. The intensities in the vessel lumen, in the basal region and in the surrounding collagen gel were extracted in order to compute the diffusive permeability (Fig. 2B-C, see methods).

In the control conditions, the diffusive permeability of 73 ± 74 nm/s was three fold larger than that of blood microvessels of ∼30 nm/s ^39^, a difference that was already reported in microvessel studies ^16^. VEGF-C stimulation moderately increased permeability of 207 ± 229 nm/s (Fig. 2B), aligning with prior reports that showed a two-fold increase in permeability in response to VEGF-C ^16,36^. In contrast, VEGF-A caused a pronounced disruption of the barrier, with a ten-fold increase in permeability to 1.1 ± 0.6 µm/s. Yet, co-stimulation with VEGF-A/C restored barrier function to levels similar to VEGF-C alone, with a permeability of 240 ± 90 nm/s. These results indicate that while VEGF-A severely compromises lymphatic barrier integrity, its deleterious effect is mitigated by VEGF-C when both growth factors are supplemented, thereby allowing enhanced sprouting without excessive degradation of barrier function.

The diffusive permeability is correlated with the quality of the barrier through the presence of voids in the tissue ^40^. To investigate the structural basis of permeability changes, we examined paracellular junctions using VE-Cad labeling in maximum intensity projections (MIPs) of confocal micrographs (Fig. 2D; see low-magnification images below). Under control conditions, the LµV exhibited continuous intercellular junctions, in agreement with previous reports ^41,42^. In contrast, the stimulation of VEGF-A induced the formation of micron-scale paracellular holes (orange arrows in Fig. 2D), which readily accounted for the onset in diffusive permeability. Such voids were not detected in response to VEGF-C or VEGF-A/C. Yet, both treatments led to the thinning of junctions, a phenotype that was enhanced with VEGF-A/C co-stimulation. To quantify the change in intercellular junction structure, we measured the full-width at half maximum (FWHM) of the VE-Cad signal, as inferred from Gaussian fits to intensity profiles (see the exemplary pink line in Fig. 2D). The FWHM was 1.71 ± 0.21 µm in control conditions. VEGF-A stimulation slightly reduced the FWHM to 1.45 ± 0.10 µm, though this difference was not statistically significant (Fig. 2E). The addition of VEGF-C alone significantly narrowed down the FWHM to 1.18 ± 0.11 µm, and junction thinning was further enhanced by VEGF-A/C co-stimulation to 1.02 ± 0.06 µm. The narrowing of the junctions under VEGF-C and VEGF-A/C in comparison to control conditions likely contributes to the moderate increase in diffusive permeability, as thinner junctions are associated with reduced apicobasal thickness, and, consequently, increased paracellular flux ^39^. These findings therefore indicate that VEGF-induced structural remodeling of intercellular junctions correlates with changes in lymphatic barrier function.

**Figure 2:**
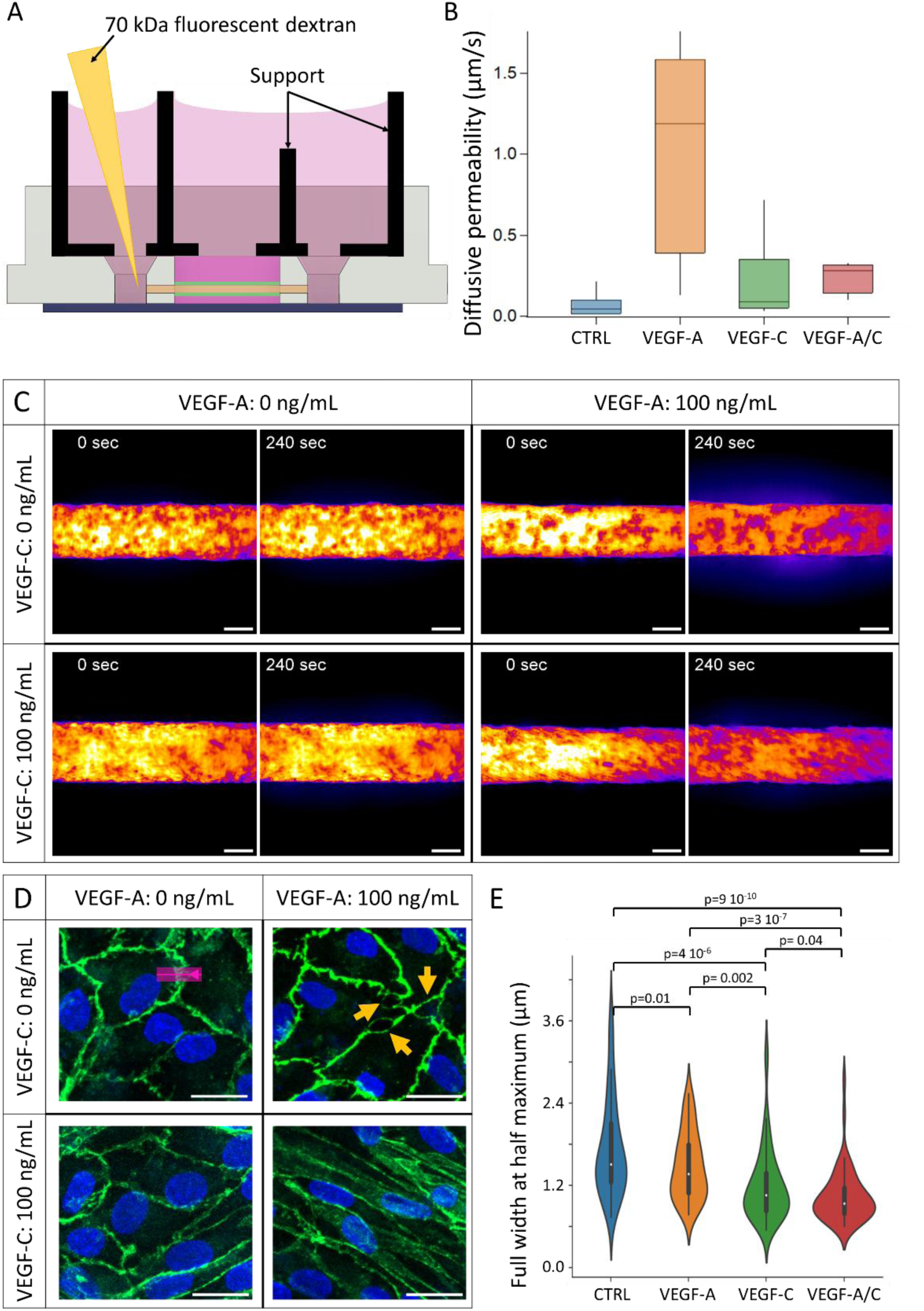
Characterization of lymphatic barrier structure and permeability. **(A)** Representation of the LµV platform and the set-up to assess diffusive permeability via injection of fluorescent dyes into the vessel lumen. **(B)** The plot presents the diffusive permeability under four conditions (control, VEGF-A, VEGF-C, VEGF-A/C co-stimulation), using n = 8–20 LµVs per condition, with measurements taken at three independent positions per vessel. **(C)** Confocal fluorescence images of LµVs filled with 70 kDa fluorescent dextran immediately after injection and after 240 s. Scale bars: 50 µm. **(D)** Maximum intensity projections (MIPs) showing nuclei and VE-Cad in blue and green, respectively. Orange arrows indicate paracellular holes. Scale bars: 20 µm. **(E)** Full width at half maximum (FWHM) of VE-Cadherin-labeled junctions across the four conditions, obtained by Gaussian fitting of intensity profiles along lines perpendicular to the junctions (example shown by the pink line in panel D).

### VEGF-A/C co-stimulation induces polarization via VEGFR-2/3 heterodimers

We next investigated the spatial organization of LECs in the tissue at the endpoint of the experiments. Using immunofluorescence confocal microscopy, we labeled intercellular junctions, filamentous actin, and DNA with VE-Cadherin, phalloidin, and Hoechst, respectively. In control samples, low magnification MIPs confirmed the continuous intercellular junction patterns of VE-Cad and revealed an actin cytoskeletal organization dominated by cortical fibers, which are characteristic of LµVs under static conditions ^16,21^ (Fig. 3A, upper left). The orientation of LECs was assessed by fitting nuclei with an ellipse in order to extract the major and minor axes, and the angle between the major axis and the vessel’s longitudinal axis (see Methods). The resulting angular distribution was approximately random (Fig; 3B), and the nuclear aspect ratio, calculated as the major to minor axis length ratio, was 1.49 ± 0.09 (Fig. 3C). VEGF-A stimulation did not alter cortical actin patterns (Fig. 3A, upper right), the nuclear aspect ratio of 1.52 ± 0.10, or the random orientation of nuclei (F = 0.2 compared to control; Fig. 3B). Note that a variance-based statistical analysis was used because the mean orientation angle is zero across all conditions due to symmetry. In contrast, VEGF-C stimulation was associated to a significant elongation of the nuclei with an aspect ratio of 1.60 ± 0.14 (p=10⁻³³), and a more pronounced alignment of the nuclei along the LµV direction (F = 0.02 compared to control; Fig. 3B-C). These trends were further enhanced after co-stimulation with VEGF-A/C (Fig. 3A, lower right), where abundant actin stress fibers were observed and nuclei alignment was peaked at zero (Fig. 3B; F < 10⁻¹⁵ compared to other conditions). The nuclei elongation also reached a maximum of 1.76 ± 0.21 (Fig. 3C). To further probe cellular polarity, we immunostained for the Golgi apparatus, a known marker of polarized cell orientation ^43,44^. Following VEGF-A/C co-stimulation, the Golgi was aligned along the nuclei main axis, while it remained randomly oriented in control conditions (Suppl Fig. S3C). Notably, this large-scale ordering of LECs was not observed in 2D cultures on either collagen gels or plastic substrates (see below and Suppl. Fig. S3A-B). Altogether, these results indicate that VEGF-A/C co-stimulation induces a phenotypic switch in LµVs, characterized by a pronounced alignment of cell nuclei specific to the vessel geometry.

The predominance of VEGFR3 is a distinguishing feature of the lymphatic endothelium, whereas VEGFR2 expression is comparable in blood and lymphatic endothelia ^45^. This status allows the specific formation of VEGFR2 homodimers, VEGFR3 homodimers, and VEGFR2/3 heterodimers in LECs ^46^. VEGFR2 homodimerization is specifically triggered by VEGF-A, whereas VEGFR3 homodimers and VEGFR2/3 heterodimers form in response to VEGF-C ^47^. To dissect the receptor contributions to polarization, we stimulated LµVs with the mutant VEGF-C156S, in which Cys156 is replaced by a Ser residue, as a selective agonist of VEGFR-3 homodimer ^48^. In contrast, mature wild-type VEGF-C triggers both homodimer and heterodimer formation. Stimulation with VEGF-C156S led to abundant and disordered actin stress fibers in fixed LµVs (Fig. 4A), associated to irregular VE-Cad patterns, an arrangement reminiscent of discontinuous *adherens* junctions with cell-cell junctions attached to the ends of stress fibers ^49^. Co-stimulation with VEGF-A/C156S reduced the formation of actin stress fibers and restored linear intercellular junctions (Fig. 4B). This result suggested co-regulatory interactions between VEGFR2 and VEGFR3 signaling pathways that are independent of heterodimer formation, similar to the co-regulation reported for the barrier function ^50^. Despite these cytoskeletal and junctional changes, the organization of nuclei in the co-stimulated tissue did not indicate polarization of the tissue, co-stimulated tissues showed no clear evidence of polarization, either by visual inspection of MIPs (Fig. 4B) or by analysis of nuclei orientation (Fig. 4C). We quantified this data by computing the difference in angular distributions between the mutant and wild-type conditions. The comparison of VEGF-C and VEGF-C156S showed a flat distribution, indicating that both growth factors similarly changed the architecture of the tissue (Fig. 4D). A flat distribution was also observed for VEGF-A/C156S *vs.* control conditions, in stark contrast to the sharply peaked distribution reported for VEGF-A/C *vs.* control (black and red datasets in Fig. 4E, respectively). Hence, our data support the conclusion that VEGFR2/VEGFR3 heterodimers, along with VEGFR2 homodimers, activated by VEGF-A/C co-stimulation, are critical drivers of lymphatic tissue polarization.

**Figure 3:**
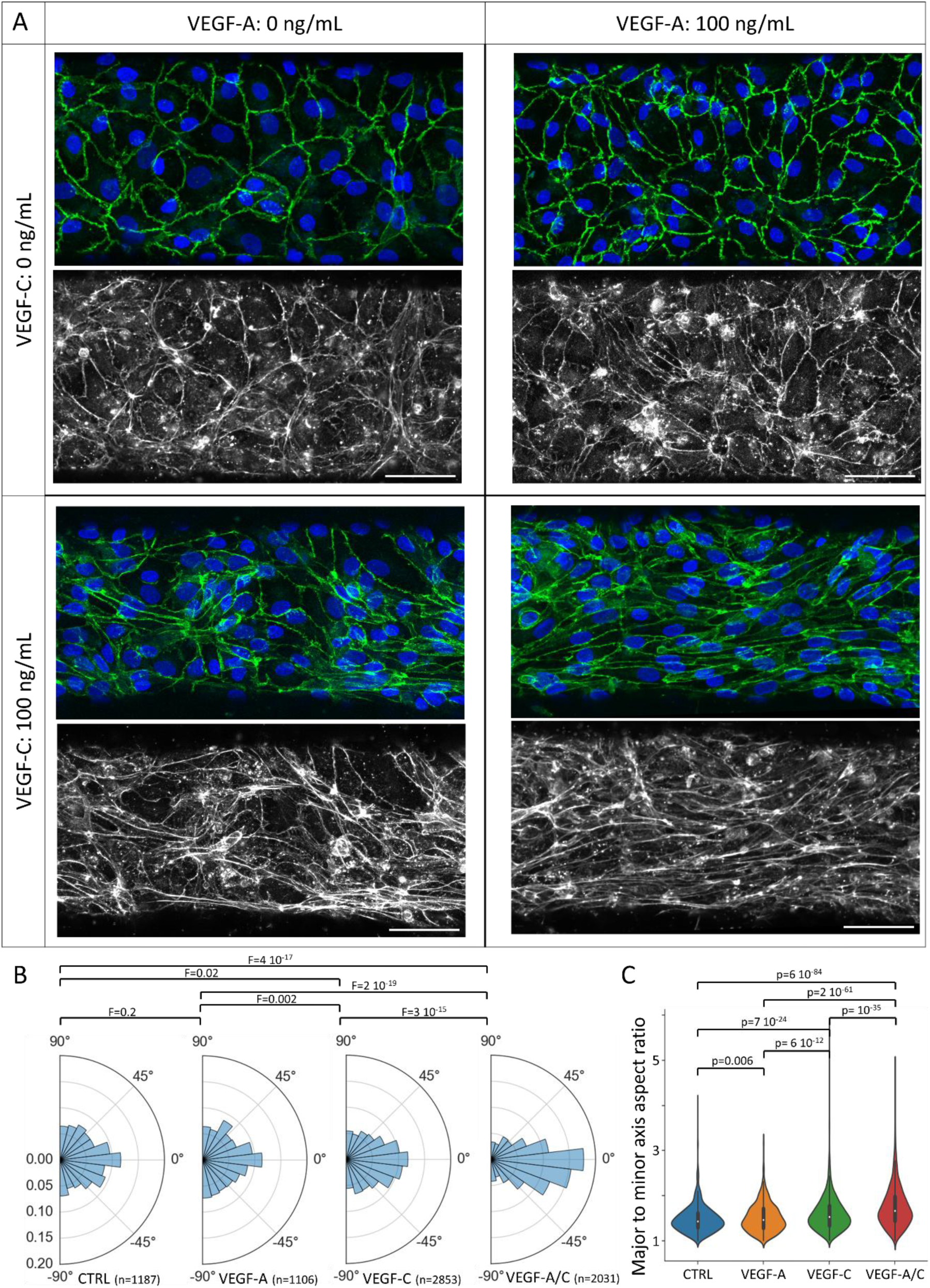
VEGF-A/C co-stimulation promotes LµV polarization. **(A)** MIPs of fixed LµVs under four experimental conditions. Upper panels show DNA (blue) and VE-cadherin (green); lower panels display F-actin stained with phalloidin. Scale bars: 50 µm. **(B)** Polar plots showing the orientation distribution of nuclei in each condition. **(C)** Quantification of nuclear aspect ratio across conditions, indicating changes in nucleus elongation associated with polarization.

**Figure 4:**
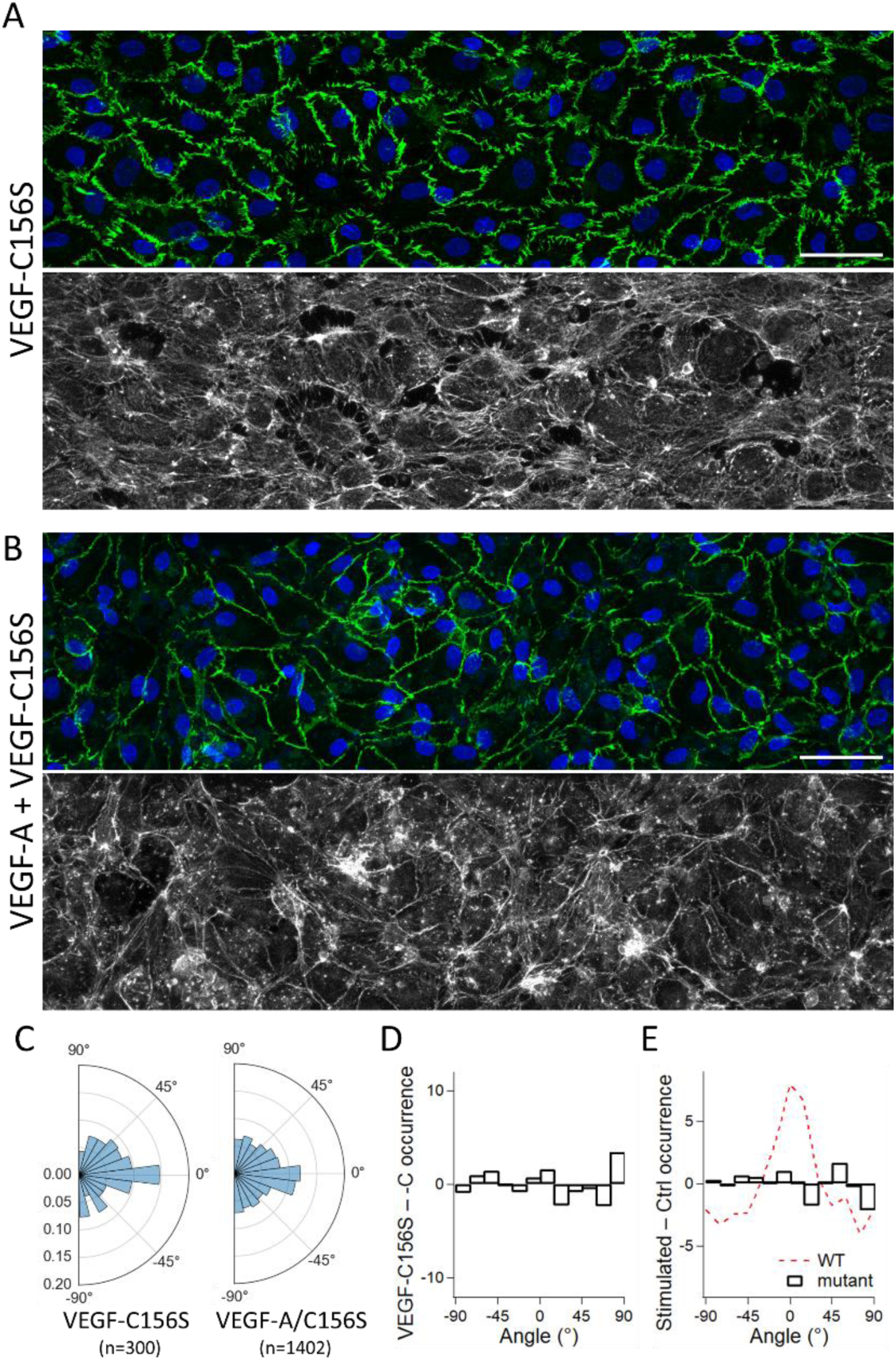
Stimulation with VEGF-C156S does not induce polarization. **(A)** MIPs of a LµV stimulated with VEGF-C156S. The upper panel shows DNA (blue) and VE-cadherin (green); the lower panel displays F-actin stained with phalloidin. **(B)** Same as (A) after VEGF-A/C156S co-stimulation. Scale bars: 50 µm. **(C)** Polar plots depicting nuclear orientation distributions upon VEGF-C156S and VEGF-A/C156S stimulation. **(D)** Differential analysis of nuclear orientation distributions comparing VEGF-C156S and wild-type VEGF-C stimulation. **(E)** Differential analysis of nuclear orientation distributions comparing VEGF-A/C156S vs. control (black histogram) and VEGF-A/C vs. control (red curve).

### Lymphatic polarization is sensitive to Rho and Src inhibition

Heterodimerization of VEGFR2 and VEGFR3 preferentially activates the PI3K/Akt signaling pathway, which is known to play a critical role in lymphangiogenesis ^51,52^. To determine whether polarization induced by VEGF-A/C co-stimulation depends on this pathway, we used SU6656, a selective inhibitor of Src family kinases ^53^, which are upstream regulators of PI3K/Akt. Co-stimulated LµVs cultured for two days in the presence of 10 µM SU6656 exhibited markedly reduced polarization, as observed in MIPs (Fig. 5A). We quantified this effect by comparing the angular distributions of nuclear orientation between co-stimulated and unstimulated conditions. The resulting difference histogram (black curve, Fig. 5A) showed only a minimal peak near 0°, indicating a lack of directional alignment. To provide a quantitative readout, we applied a Gaussian fit with an offset to the angle distribution (red curve, Fig. 5A), using a two-parameter model (amplitude and width), constrained such that the integral was zero (see Methods). he fit yielded a low amplitude (2.1 ± 2.3) and a poor goodness-of-fit score (92.3), reflecting the absence of a well-defined directional bias. In contrast, the same analysis performed on VEGF-A/C co-stimulated samples without SU6656 showed a higher amplitude (10.4 ± 0.9) and a substantially better fit (goodness-of-fit score: 11.2; Suppl. Fig. S4A). These findings indicate that SU6656 disrupts tissue polarization by interfering with PI3K/Akt-mediated mechanotransduction.

We next investigated the role of the Mitogen-Activated Protein Kinase (MAPK) pathway, another mechanotransduction cascade active in endothelial cells. Specifically, we focused on the p38 and ERK1/2 branches, both of which are known to be activated by shear stress ^54^. To assess the contribution of p38 MAPK, we used SB203580, a selective inhibitor previously shown to reduce lymphangiogenesis in cancer models ^55^. Treatment with SB203580 did not impair LµV polarization, as indicated by MIPs (Fig. 5B). Gaussian fitting of the difference in nuclear orientation distribution yielded an amplitude of 10.5 ± 2.5 that was comparable to untreated controls, thus confirming the occurrence of polarization. We then examined the role of ERK1/2 signaling using PD98059, an inhibitor known to block the rapid activation of ERK1/2 by VEGF-A ^51^. Similar to p38 inhibition, PD98059 did not significantly affect LµV polarization. The Gaussian fit returned a goodness-of-fit score of 47.9 and an amplitude of 7.2 ± 1.7 (Fig. 5C), indicating preserved polarization. Together, these results suggest that inhibition of either p38 or ERK1/2 MAPK signaling does not significantly disrupt LµV polarization in response to VEGF-A/C co-stimulation.

Given that VEGF-A/C triggers the formation of actin stress fibers (Fig. 3A), we asked whether targeting ROCK — a kinase activated by RhoA that regulates myosin motor activity and actin stress fiber formation — with the inhibitor Y27632 would interfere with tissue polarization. By reducing the formation of stress fibers, this inhibitor has been shown to contribute to the stabilization the lymphatic barrier function when challenged by inflammatory factors such as histamine ^56^. After two days of culture with Y27632, the polarization of LµVs was essentially blocked in MIPs (Fig. 5D). The difference of nuclei angle distribution was roughly random (right panel in Fig. 5D), as confirmed by the largest score for the goodness-to-fit (118.6) and the negligible amplitude of the signal (4.0 ± 3.0). Taken together, these results demonstrate that VEGF-A/C-induced polarization of LµVs critically depends on the PI3K/Akt mechanotransduction pathway and ROCK kinase activity.

Notably, inhibition of polarization by SU6656 or Y27632 did not prevent the formation of lymphangiogenic sprouts, as confirmed by both brightfield and confocal microscopy (Fig. 5E and Supplementary Fig. S4B). To test the converse, *i.e.*, whether blocking sprouting would affect polarization, we cultured LµVs in a cross-linked collagen matrix (see Methods), a condition that restricts LEC migration and significantly suppresses sprouting (Supplementary Fig. S4C-D). Despite the inhibition of sprouting, VEGF-A/C-induced polarization remained evident (Supplementary Fig. S4E). These findings suggest that lymphangiogenic sprouting and tissue polarization are mechanistically distinct processes. Although they occur in parallel under native collagen conditions, they can be uncoupled by altering the biophysical properties of the extracellular matrix.

**Figure 5:**
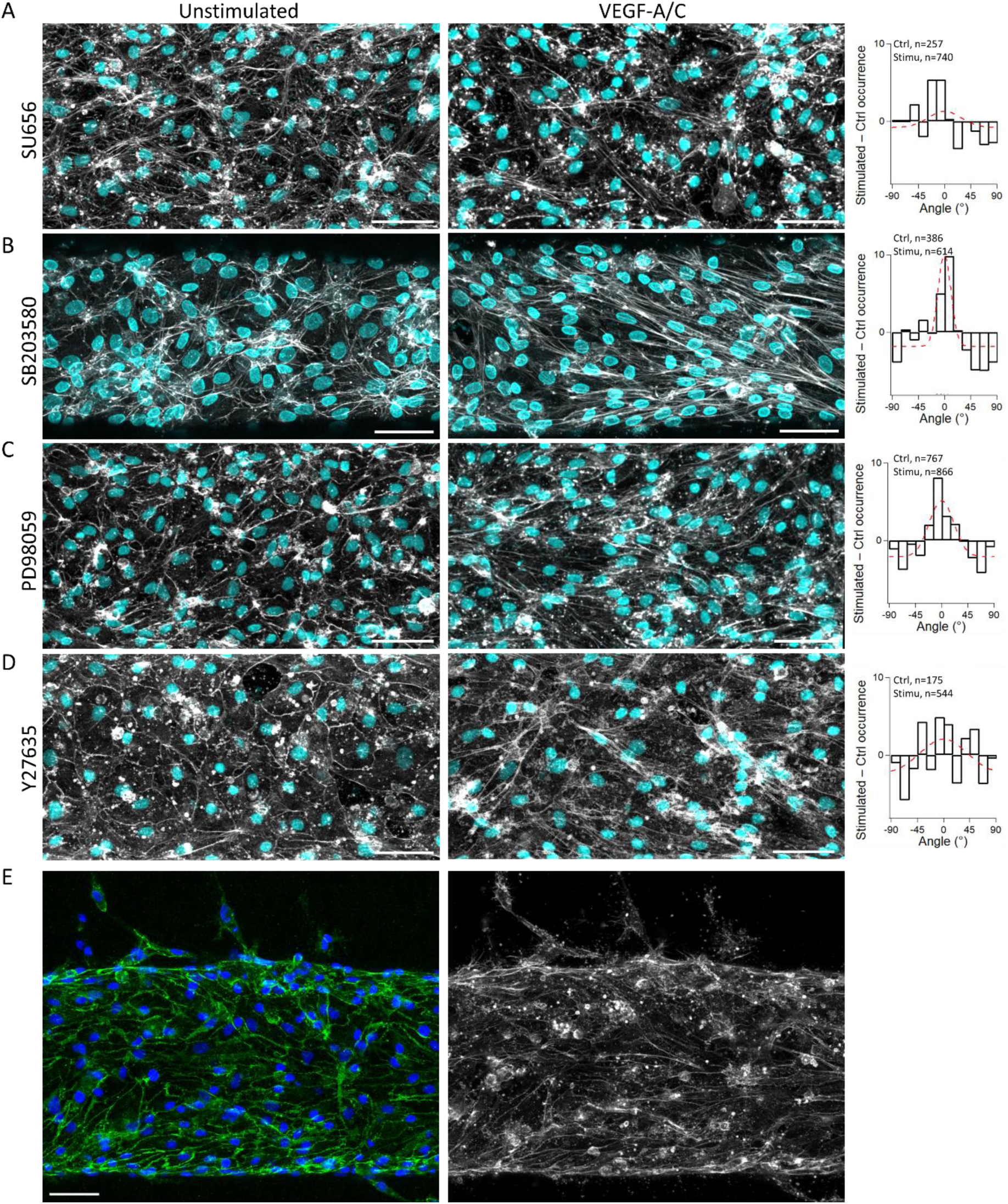
Src and ROCK inhibition disrupt LµV polarization. **(A)** MIPs of LµVs treated SU6656 (Src inhibitor) under stimulated or unstimulated conditions The histogram on the right shows the difference in nuclear orientation distributions between the two conditions, along with a Gaussian fit (see Methods). **(B-D)** Same analysis as in (A), using SB203580 (p38 MAPK inhibitor), PD98059 (MEK/ERK1/2 inhibitor), and Y27632 (ROCK inhibitor), respectively. **(E)** MIP of an LµV treated with Y27632 and co-stimulated with VEGF-A/C. DNA (blue), VE-cadherin (green), and F-actin (gray, phalloidin stain) are shown. All inhibitors were used at a concentration of 10 µM. Scale bars: 50 µm.

### Anisotropic durotaxis as a tentative mechanism driving tissue polarization

While we investigated the roles of specific receptors and mechanotransduction pathways in LµV polarization, the physical mechanism by which a biochemical signal is translated into large-scale tissue reorganization remains unresolved. Given that VEGF-C has been shown to enhance LEC migration, in particular by activation of PI3K ^35^, we first focused LEC motility in a simplified 2D setting to decouple motion from the geometric complexity of vessels (see Methods). Confluent LEC monolayers cultured on collagen gels were stimulated overnight with VEGF-A, VEGF-C, VEGF-A/C, or VEGF-A/C156S (Fig. 6A), and individual cell trajectories were tracked for 8 hours at 10-minute intervals. We quantified motility using mean square displacement (MSD; Fig. 6B) and particle image velocimetry (PIV; Supplementary Fig. S5A). After four hours, MSD values normalized to control increased by 15%, 29%, 120%, and 289% following stimulation with VEGF-A/C156S, VEGF-A, VEGF-C, and VEGF-A/C, respectively. These results demonstrate a synergistic effect of VEGF-A/C on motility, which was absent with the VEGF-C156S mutant.

In control, VEGF-A, and VEGF-A/C156S conditions, MSD increased linearly with time (e.g., green fit in Fig. 6C), consistent with purely diffusive, random motion. From the slope, we inferred effective diffusion coefficients of 60 ± 1, 76 ± 1, and 68 ± 1 × 10⁻³ µm²/min, respectively. In contrast, VEGF-C and VEGF-A/C stimulation produced MSD curves that deviated from linearity and fit well to parabolic trends (Fig. 6C), indicating drift-diffusion behavior, *i.e.*, motion with directional persistence on the 10-minute timescale. The average velocities under VEGF-C and VEGF-A/C were 32 ± 2 and 45 ± 3 × 10⁻³ µm/min, respectively. A diffusive component was still present, with diffusion coefficients of 76 ± 5 and 118 ± 5 × 10⁻³ µm²/min, respectively. Notably, while the baseline diffusive motion was comparable between VEGF-A and VEGF-C, only VEGF-C (and more strongly VEGF-A/C) induced persistent, directed migration. PIV analysis further revealed that cell displacements were not uniformly distributed but instead formed dynamic, large-scale velocity domains across the field of view (Supplementary Fig. S5), a hallmark of collective migration ^57,58^. Together, these results indicate that VEGF-A/C co-stimulation promotes both increased motility and enhanced collective coordination among LECs, pointing to a mechanistic link between receptor signaling and tissue-level organization.

We next investigated how LµV geometry influences the polarization process by measuring vessel diameter immediately after growth factor addition and again after 16 hours of stimulation. By calculating the ratio of these measurements, we found that VEGF-A/C co-stimulation led to vessel retraction, whereas control and VEGF-C treatments resulted in slight expansion (Fig. 6D). This retraction suggests the generation of tensile forces between adjacent LECs. To understand how such forces impact tissue deformation, we conducted finite element simulations modeling the lymphatic vessel as a homogeneous elastic solid subjected to prestress along both the longitudinal and equatorial axes (see Methods). Simulation results revealed that tensile forces predominantly induced deformation in the equatorial plane, with minimal displacement along the longitudinal axis (Fig. 6E). This behavior can be attributed to the inherent curvature of the vessel wall, which redirects tensile forces inward toward the centerline, producing circumferential contraction. The asymmetric mechanical response, compliant in the equatorial direction and stiff longitudinally, combined with the enhanced motility observed under VEGF-A/C stimulation, supports a polarization mechanism reminiscent of durotaxis ^59^. In particular, it resembles directional migration along the stiff axis of substrates with anisotropic mechanical properties, such as micropillar arrays with elliptical cross-sections ^60^ or aligned liquid crystalline polymer networks ^61^. However, unlike these engineered systems, we propose that in microvessel models, stiffness anisotropy arises intrinsically from the tubular geometry of the LEC monolayer. We term this mechanism anisotropic durotaxis. Notably, Src has been shown to regulate cell contractility ^62^, linking VEGF-A/C-mediated polarization to geometry-guided migration through a Src-dependent anisotropic durotaxis mechanism.

**Figure 6:**
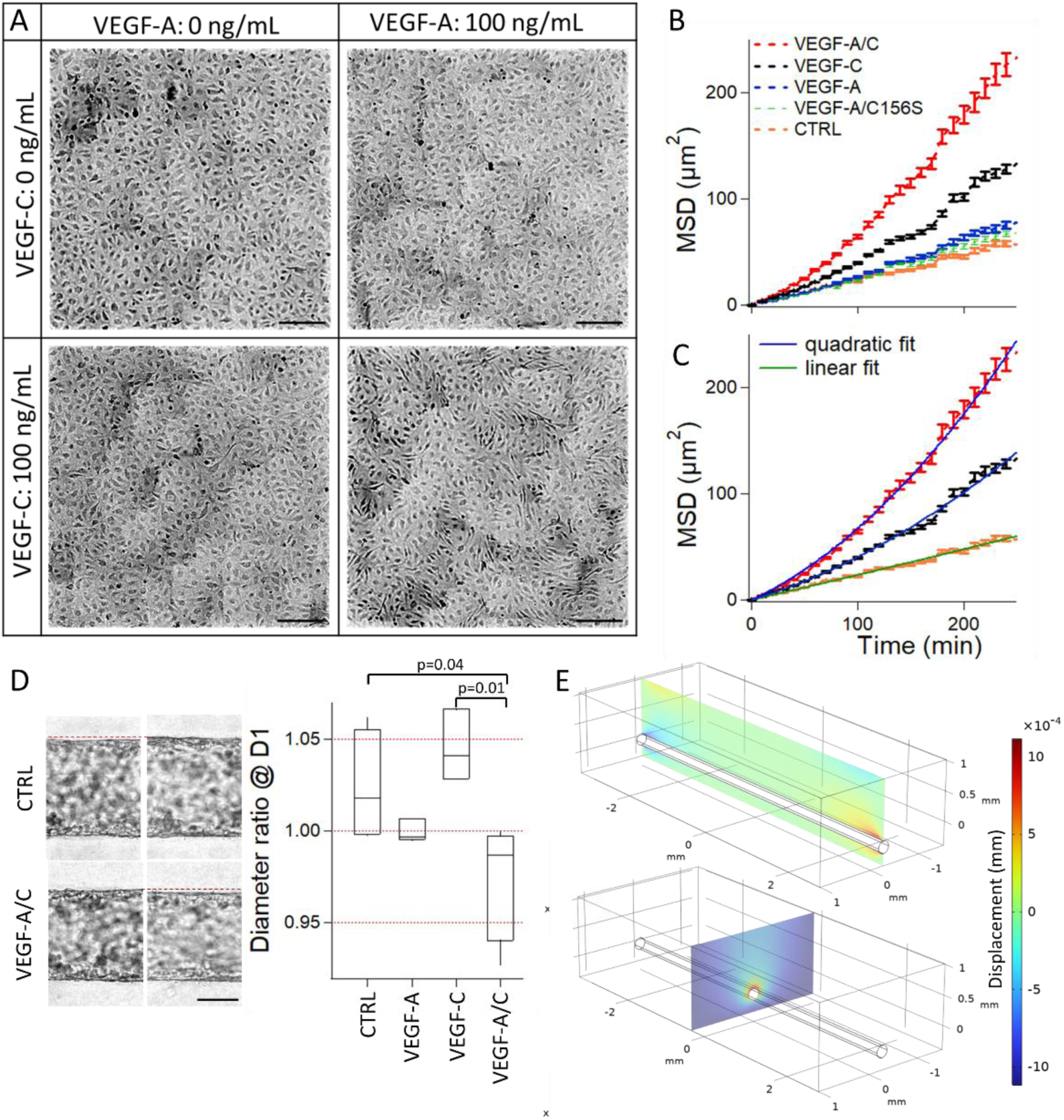
Mechanism of LµV polarization. **(A)** Bright field images of LECs cultured on flat collagen gels after 24 hours of stimulation under different conditions. Scale bars: 20 µm. **(B)** Mean square displacement (MSD) plotted as a function of time for five experimental conditions (see legend). **(C)** Selected MSD curves from (B) fitted with either linear (diffusive, blue line) or quadratic (drift-diffusion, green line) models. **(D)** Ratio of LµV diameter measured at 16 hours relative to 3 hours, based on contour analysis of brightfield images. Representative images are shown with red dashed lines as a visual guide to diameter changes. Scale bar: 100 µm. **(E)** Finite element simulations showing tissue deformation resulting from isotropic prestress (i.e., contraction forces between adjacent cells) in the longitudinal and equatorial directions. Simulations reveal minimal longitudinal deformation (top panel, green) and maximal equatorial deformation (bottom panel, red), as indicated by the color scale.

## Conclusion

In this study, we developed a tissue engineering approach to fabricate a lymphatic vessel-on-a-chip within a collagen gel matrix. This platform enabled us to examine the response of LµVs to VEGF-A and VEGF-C—key endothelial growth factors—by integrating data from bright-field and confocal immunofluorescence microscopy. Co-stimulation with VEGF-A/C induced large-scale tissue polarization, a phenomenon not previously described in the literature to our knowledge. This effect depends on the formation of VEGFR2–VEGFR3 heterodimers. It is impaired using inhibitors directed to either PI3K/Akt, a signaling cascade triggered by VEGFR2–VEGFR3 heterodimers, or ROCK. Mechanistically, we demonstrate a synergetic enhancement of cell motility under VEGF-A/C co-stimulation, and propose that polarization results from directed cell migration guided by anisotropic mechanical response of the vessel’s cylindrical geometry, stiff in the longitudinal axis and soft in the equatorial plane. Notably, lymphatic vessels undergo dynamic remodeling during development, particularly in response to VEGF-C stimulation. While fluid shear stress has been suggested to contribute to these reorganizations ^10^, its notoriously weak strength in lymphatic vasculatures ^64^ has not lent support to this hypothesis. Our data prove that polarization can be induced by biochemical cues. However, unlike *in vivo* observations equatorial polarization ^10–12^, we report an axial alignment. Given the role spatial gradients in VEGF-induced lymphangiogenesis ^34^, we suggest to investigate how spatially-resolved VEGF-A/C co-stimulation patterns influence the polarization dynamics, and in turn contribute to the formation of specialized lymphatic structures both *in vitro* and *in vivo*.

We also demonstrated that VEGF-A/C co-stimulation synergistically promotes sprouting lymphangiogenesis without significantly compromising lymphatic barrier integrity. Given that maintaining a functional lymphatic barrier is essential for proper fluid drainage and immune surveillance, this result underscores the importance of tightly regulating the balance between growth-promoting signals and vascular stability. Excessive or uncoordinated stimulation may drive uncontrolled remodeling at the expense of vessel integrity, potentially impairing lymphatic function. Thus, our data hint to the key role of growth factor signaling coordination to drive lymphangiogenesis and preserve barrier properties, a consideration that may be relevant in therapeutic or pathological contexts where VEGF levels can be elevated ^63^. Another intriguing observation is that intradermal injection of VEGF-A or VEGF-C in mice promotes the formation of zipper-like LEC junctions ^41^. In our *in vitro* system, co-stimulation led to reorganization of VE-cadherin junctions into thin, linear structures reminiscent of zipper junctions. However, unlike the *in vivo* context, where the transition from button-to zipper-like junctions correlates with enhanced barrier function ^65^, we observe that these linear junctions are associated to a modest destabilization of the lymphatic barrier. Although reproducing the button-to-zipper junction transition remains, to our knowledge, a challenge in LµV platforms, we argue that their controllable molecular and mechanical environment offers a valuable tool for dissecting the key mechanisms underlying lymphatic maturation and junctional remodeling.

## Material and Methods

### Cell culture and reagents

Human dermal lymphatic endothelial cells (HDLECLot 483Z001.2, PromoCell) were cultured in Microvascular Endothelial Cell Growth Medium-2 BulletKit (EGM-2, Lonza) in standard tissue culture incubators at 37°C, 95% humidity, and 5% CO2. Cells were used at passage 5 and 6. Stock solutions of VEGF-A, VEGF-C (ThermoFischer), VEGF-C156S (R&D systems) were prepared in sterile phosphate buffer saline (PBS, Sigma-Aldrich), aliquoted, and stored at - 20°C until further use. Supplemented media with VEGFs was prepared by diluting stock solution in fresh Lonza medium at 100 ng/mL. Medium containing both of VEGF-A and VEGF-C was supplemented by both growth factors at 100 ng/mL each. PD98059, SU6656, SB203580 (Sigma Aldrich) stock solutions were prepared in sterile DMSO, used at a concentration of 10 µM in medium. Y27632 (Sigma Aldrich) was diluted in water and used at a concentration of 10 µM. All inhibitors stock solution were stored at -20°C, and supplemented media was prepared by thawing and diluting inhibitor stock solutions at the desired concentration.

### Lymphatic micro-vessels fabrication (LµV) & growth factor stimulation

LµV were prepared as described in a previous report ^37^. Silicone chips were treated with oxygen plasma for 30 s and placed in a vacuum chamber with 50 µL of 3-aminopropyl-triethoxysilane (Sigma Aldrich) for 30 min. Acupuncture needles (No.3, J type; Seirin) were coated with 1% (w/v) bovine serum albumin (BSA) solution for 15 min and air dried. Under culture hood, silanized chips were treated with 60 µL of 2.5% glutaraldehyde (GA; ThermoScientific), then thoroughly rinsed with sterile DI water and air dried. A collagen solution was prepared on ice by mixing Cellmatrix® Type I-A Collagen (Nitta Gelatin), 10x Hanks’ Balanced Salt Solution (Sigma Aldrich) and 10x collagen buffer reagent (Sodium hydroxyde 50mM, Sodium hydrogen carbonate 260 mM, HEPES 200 mM, Sigma Aldrich) at a 8:1:1 volume ratio with a final collagen concentration of 2.4 mg/mL. 40 µL of this solution was poured in the vessel chip before inserting the BSA-coated needle, then incubated at 37 °C for 15 min to allow reticulation. Chips were stored in sterile PBS at 4 °C overnight to prevent bubble formation. Needles were withdrawn and channels coated with a 10 µg/mL solution of Human Fibronectin (MilliPore) for 30 min. At this stage, collagen scaffolds can be cross-linked by soaking a 1% para-formaldehyde (PFA, Sigma Aldrich) solution in the chip for 30 minutes, and then extensively rinsing chips for 3 days. For seeding, confluent HDLEC cultures were re-suspended in EGM-2 MV media supplemented with 3% (m/v) 500 kDa Dextran (Sigma Aldrich) at a concentration of 10 × 10^6^ cells/mL. Cells were seeded by sequential loading of 5 µL of the cell suspension in each reservoir, allowing cells to flow and attach into the channel. In between loadings, chips were flipped upside down, and incubated at 37°C for 10 min, before reiterating seeding until full coverage of the vessel was achieved. Chips were eventually filled with 1 mL of fresh medium, and stored at the incubator for two hours. Then, media was changed according to the tested condition (VEGFs, inhibitors), and kept at the incubator up to 3 days. Chips were imaged 2 times a day with a brightfield optical microscope (Olympus) equipped with a 20X air objective.

### Permeability assay

After 3 days of culture, LµV were retrieved from the incubator and their barrier function individually assessed, as described in ref. ^37^. Briefly, chips were placed in an aluminum support, and a dedicated 3D-printed support was screwed on top of the chip, allowing a separation of the main and side reservoirs. Once filled with warm culture media, the assembly was placed under confocal microscope (Leica) at 10X and focus was made in the middle of the vessel. A solution of 70 kDa Dextran-FITC (20mg/mL, Sigma Aldrich) was added to one of the lateral inlets and flowed into the lumen. Fluorophore diffusion into the collagen was imaged at 3 different positions every 10-20 s for 5 min, and the resulting timelapse was processed using a custom ImageJ macro to retrieve diffusive permeability values (*L_D_*) for each position, as defined by the following expression:

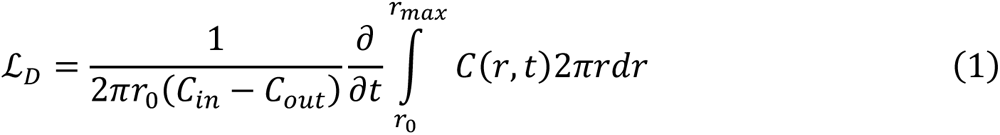

with *r*_0_ the vessel radius, *C_in_*, *C_out_*, and *C(r, t)* the apical, basal, and bulk collagen intensities, respectively. The averaged value was used as sample value. Subsequently, vessels were fixed by adding 500 µL of 1% PFA solution for 30 min, rinsed with PBS, and stored at 4°C until further characterization.

### Immunostaining and imaging of LµV

Fixed samples were permeabilized with 1 mL of 0.5% Triton-100X (Sigma-Aldrich) in PBS for 10 min, and blocked with BSA (10 mg/mL, Sigma-Aldrich) overnight at 4 °C. Primary antibody detecting GM130 (Rabbit, 1:200; Cell Signaling) were incubated in a 1% BSA solution overnight at 4°C in dark. After washing, cells were incubated for 2h with secondary antibodies (Anti-rabbit, 1:400, Jackson), Phalloidin (1:1000, Cell Signaling) and DAPI (1:1000, Sigma Aldrich) at room temperature in dark. Conjugated antibody detecting VE-cadherin (1:200; Mouse, Santa Cruz coupled with 488 Anti-mouse, Jackson) was incubated in 1% BSA solution for 2h. After washing, DAPI (1:1000, Sigma Aldrich) was incubated for 10 min. Labeled samples were washed and stored at 4°C in dark before imaging. Confocal images were acquired using the LSM900 confocal microscope (Zeiss) with a 25X water immersion objective (LD LCI Plan-Apochromat 25x, Zeiss) or using the LSM700 confocal microscope (Zeiss) with 20X objective (LD Plan-Neofluar 20X, Zeiss). Confocal stacks were recorded over a depth of 20 to 30 µm with confocal slices separated by 1 µm starting from the bottom of the vessel, and maximum intensity projections (MIPs) were computed as final readout.

### 2D live imaging assays

2D-coated 12-well plates were prepared from standard 12-well plates (Corning). Plates were put through oxygen plasma (10s, 100%), and UV-sterilization (5 min). A collagen type I-A solution was prepared as mentioned, and each well was coated with 200 µL of collagen. After 10 min of reticulation, sterile PBS was added to each well, and plates were stored at 4 °C until further use. The day of the experiment, LEC were passed and seeded onto the collagen-coated wells at a concentration of 0.1 × 10^6^cells/mL, and incubated until confluency was reached. Media was then replaced by VEGF-supplemented media according to the tested conditions, and incubated overnight. The next day, the well plate was placed under a motorized microscope (Observer 7, Zeiss) and brightfield images tiles of each well were taken every 10 min for 9 hours.

### Numerical simulations

We performed finite element simulations with the mechanics module of COMSOL. We reproduced the geometry of the vessel chip, and parametrized the elasticity of collagen gel to 250 Pa based on our measurements ^66^ (note that there is no loss of generality due to linear physics). The cell monolayer was modeled as an homogeneous material of 3 µm in thickness, its elasticity was equal to the Young modulus of individual LECs of 2 kPa ^67,68^. Finally, we assumed a pre-stress in the simulation resulting from contractile forces of 500 Pa ^69^ homogeneous in the longitudinal and equatorial axis.

### Image analysis

FIJI was used for image analysis procedures ^70^. Contrast of brightfield images was enhanced using the *Remove background* (Rolling ball radius: 50px), *Enhance contrast* (Saturated pixels 0.35%; Normalization) functions. For LµV contour length measurements and lymphangiogeneic sprout quantification, the LµV and collagen matrix were segmented using the Trainable Weka Segmentation plugin for FIJI ^71^. Briefly, a custom classifier was created by training the plugin on multiple images for each VEGF conditions, and applied in batch to all the images. Relative contour length at day 3 was calculated from the normalized contour length 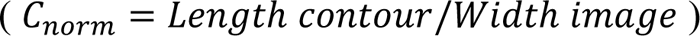 and compared to the value at day 0 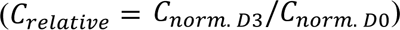.

For nuclei angle measurements, the StarDist plugin for FIJI ^72^ was used to segment the nuclei channel and retrieve individual nuclei characteristics (number of nuclei, angle, aspect ratio). For 2D assays, brightfield images were first segmented using the Trainable Weka Segmentation plugin, and cells were then tracked using the TrackMate plugin for FIJI ^73,74^. Tracks were exported and the mean square displacement for each condition was calculated using the CelltrackR package from the MotilityLab webpage ^75^.

### Fits, graphs and statistics

Graphs and fits were obtained using Igor Pro 5 or Python’s seaborn package ^76^. The normalized Gaussian function used to fit the data in Fig. 5 was defined as

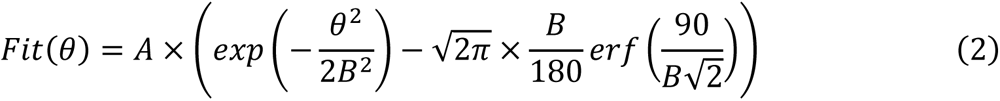

with *A* and *B* the fitting parameters, and the FWHM equal to 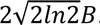. For MSD analysis, we used the diffusion and convection solution

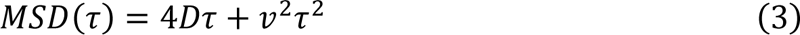

with *D* the diffusion coefficient, *v* the velocity, and *τ* the time lag. Statistical significance was determined across our data using a Student’s t-test under a normal distribution assumption. p < 0.05 was the threshold set for statistical significance. For angle distributions, we performed f-tests for variance-based statistical significance.

## Acknowledgments

J.E. thank the CNRS international program for his PhD fellowship. B.A. acknowledge the JSPS for a postdoctoral fellowship.

## Supplementary material

**Supplementary Fig. S1:**
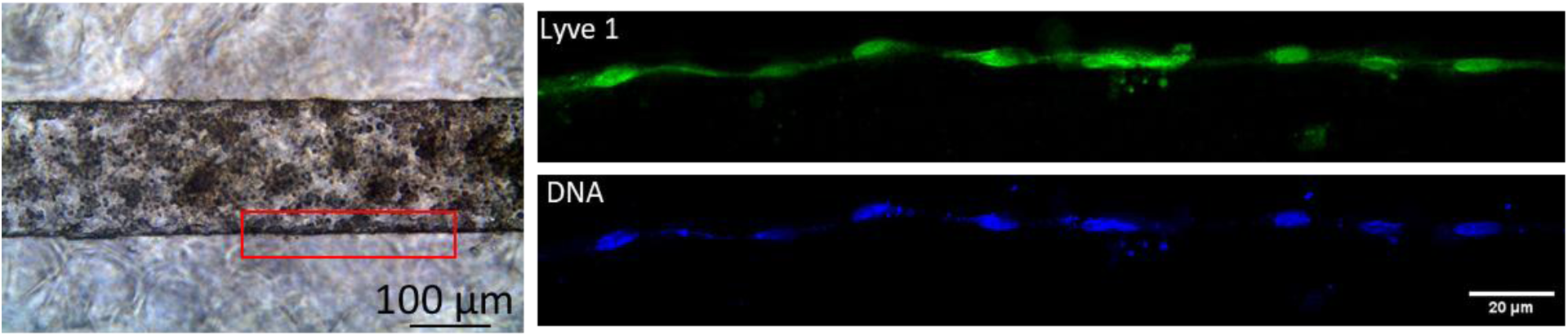
Immunoconfocal stack performed on the side wall of a LµV labeling DNA and Lyve-1.

**Supplementary Fig. S2:**
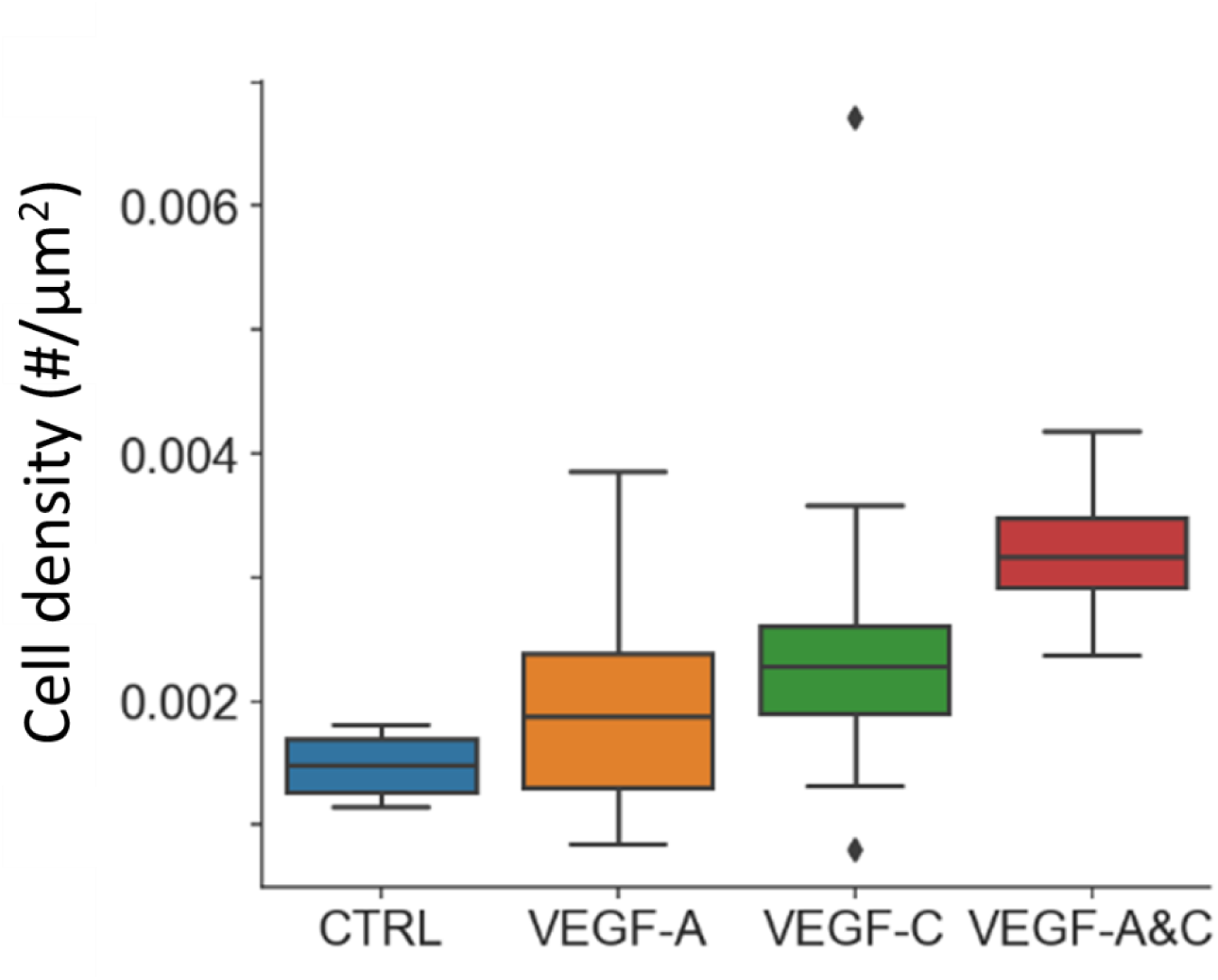
Density of nuclei in LµV after three days of culture. The counting is performed on MIPs of 20 µm in thickness. We use 5 independent chips for each condition, and enumerate the number of nuclei in three images for each LµV.

**Supplementary Fig. S3:**
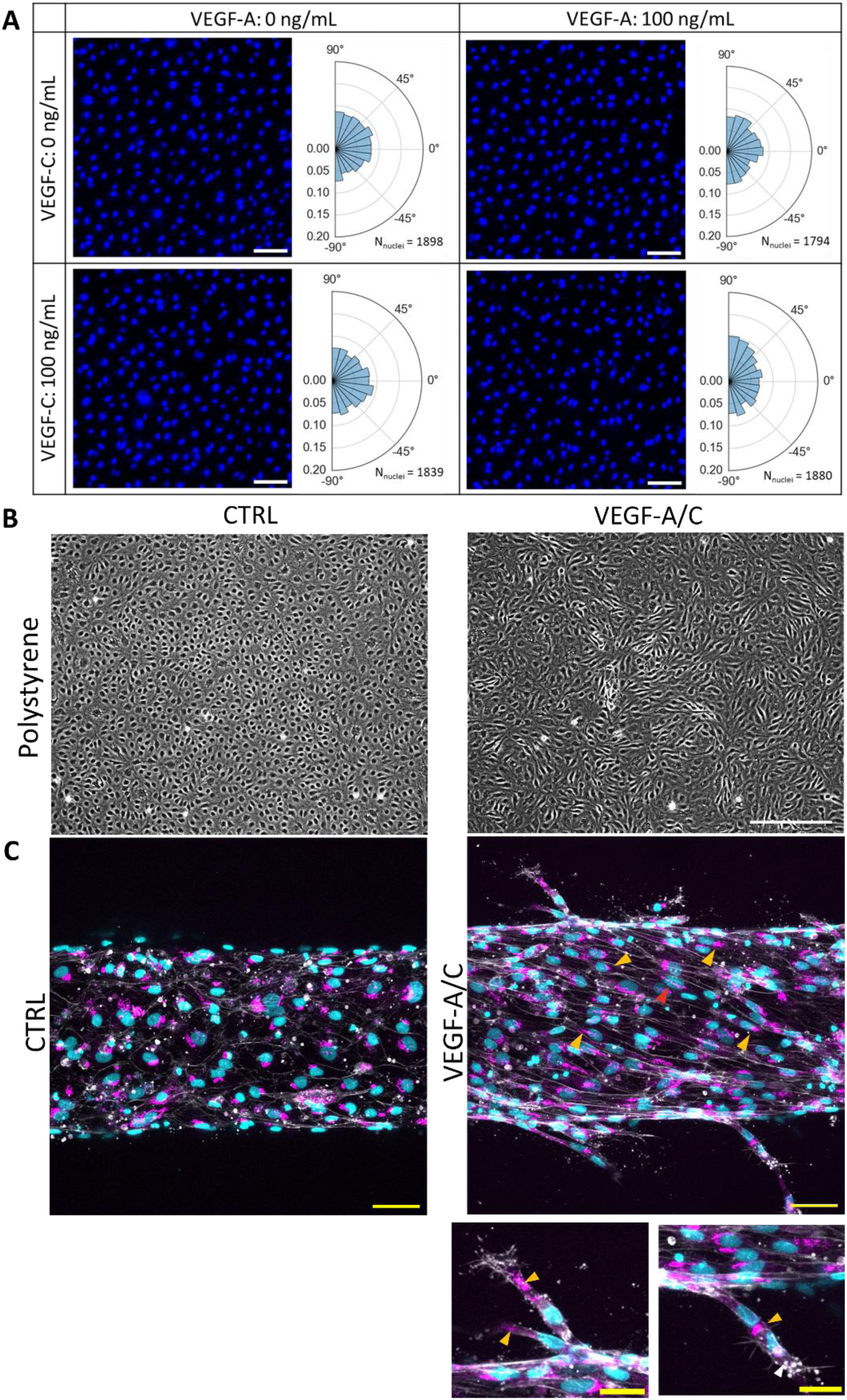
**(A)** Fluorescence micrographs of LEC nucleic labeled with Hoechst grown on flat collagen gels in control, VEGF-A, VEGF-C, and VEGF-A/C conditions. Histogram of nuclei orientation are appended to the images. Scale bars: 20 µm. **(B)** Wide-field microscopy images of LECs cultured on conventional Petri dishes under control and co-stimulated conditions. Scale bars: 20 µm. **(C)** MIPs of LµVs stained with Hoechst (blue), phalloidin (gray), and an anti-Golgi apparatus antibody (pink). Orange arrowheads indicate instances where the Golgi apparatus is aligned with the nucleus’s major axis, typically observed in sprouts or polarized tissues. The red arrowhead highlights a Golgi apparatus encircling the nucleus along its main axis orientation. Scale bars: 50 µm in low mag images and 25 µm in zoom-ins.

**Supplementary Fig. S4:**
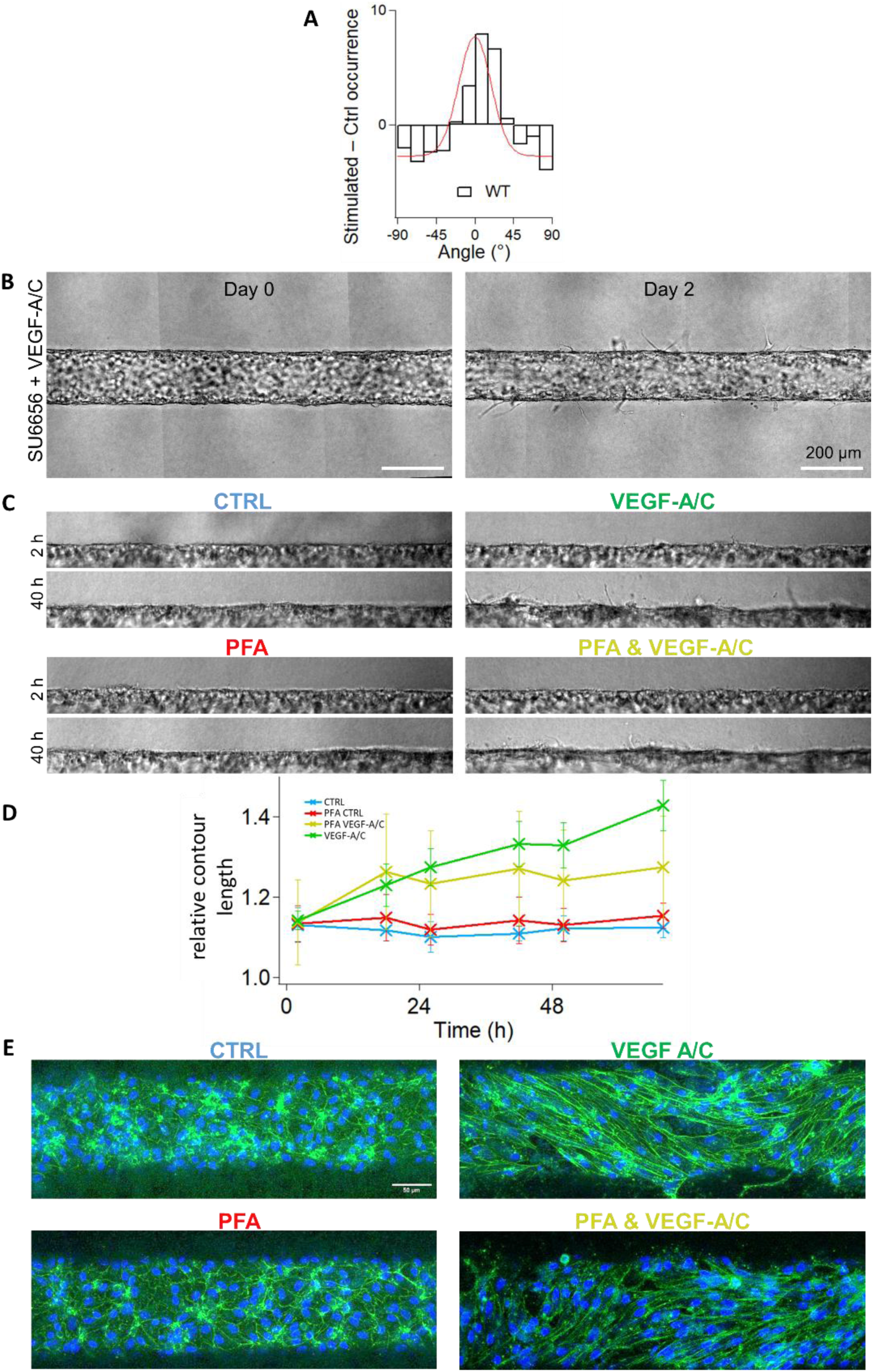
**(A)** Differential analysis of nuclear orientation distributions comparing VEGF-A/C *vs*. control (black histogram) and its fit (red curve). **(B)** Wide-field micrographs of LµVs treated with SU6656 just after the fabrication and two days later. Sprouts are detected after drug treatment. **(C)** Wide-field micrographs of co-stimulated LµVs fabricated in native collagen gels (upper panels) or cross-linked collagen gels (lower panels). **(D)** The formation of sprouts is inhibited in cross-linked matrices, in which the contour length increases slightly during the first 16 hours of the experiment and remains constant after (yellow curve). The relative contour length increases in native gels (green curve). **(E)** MIPs of LµVs in control or co-stimulated conditions after two days of culture. Labels: DNA in blue and phalloidin in green.

**Supplementary Fig. S5:**
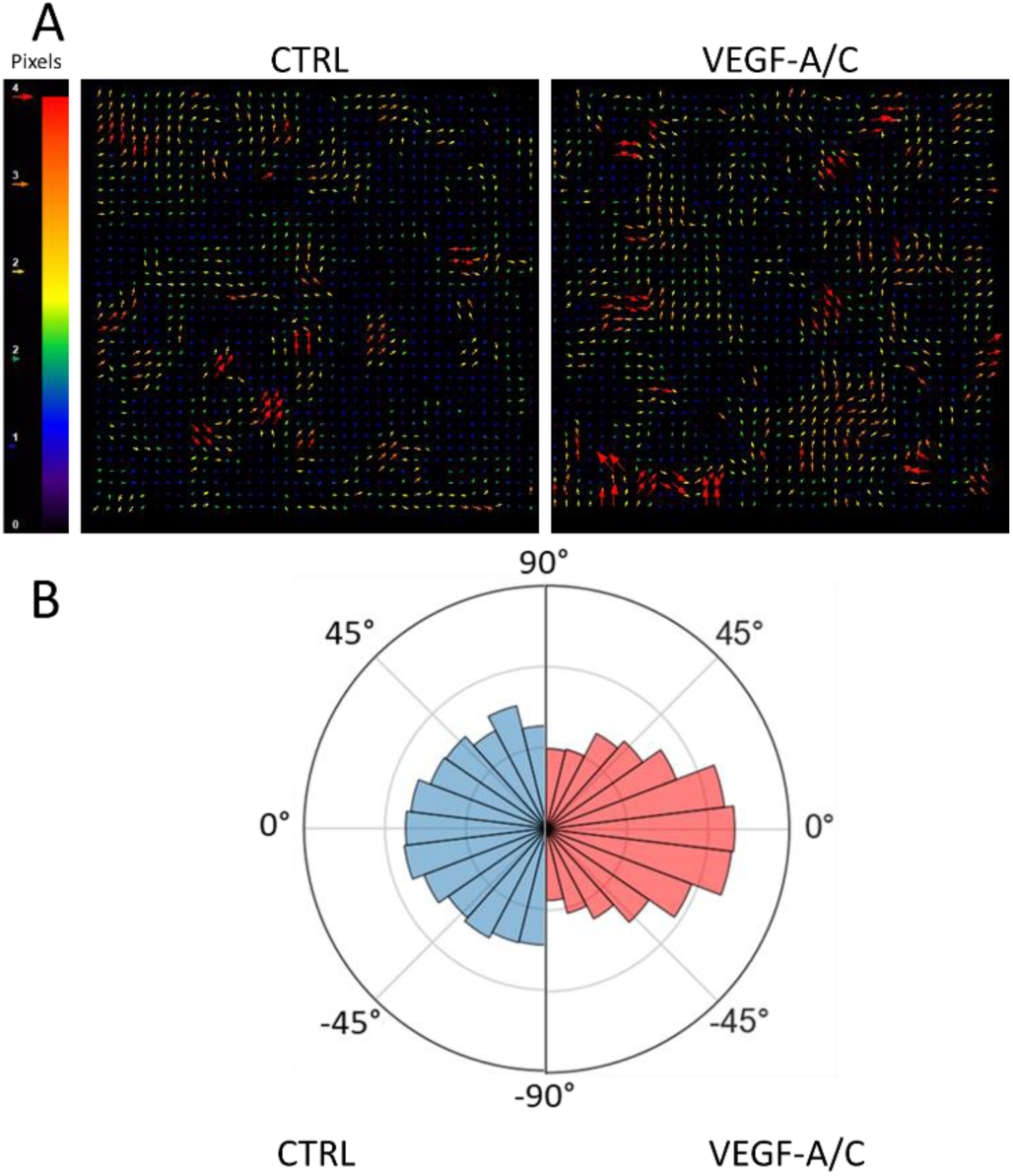
**(A)** PIV maps in control and co-stimulated conditions (left and right panels, respectively). **(B)** The polar plots represent the difference of angle between two nuclei distant of less than 50 µm in 2D cell culture experiments. The angle is random in control conditions but correlated (i.e., frequently close to 0) under co-stimulated conditions.

## Bibliography

1. Aspelund, A., Robciuc, M.R., Karaman, S., Makinen, T., and Alitalo, K. (2016). Lymphatic System in Cardiovascular Medicine. Circ. Res. 118, 515–530. 10.1161/CIRCRESAHA.115.306544.

2. Petrova, T.V., and Koh, G.Y. (2018). Organ-specific lymphatic vasculature: From development to pathophysiology. J. Exp. Med. 215, 35–49. 10.1084/jem.20171868.

3. Zhang, F., Zarkada, G., Yi, S., and Eichmann, A. (2020). Lymphatic Endothelial Cell Junctions: Molecular Regulation in Physiology and Diseases. Front. Physiol. 11, 509. 10.3389/fphys.2020.00509.

4. Suami, H., and Scaglioni, M.F. (2018). Anatomy of the Lymphatic System and the Lymphosome Concept with Reference to Lymphedema. Semin. Plast. Surg. 32, 5–11. 10.1055/s-0038-1635118.

5. Moore, J.E., and Bertram, C.D. (2018). Lymphatic System Flows. Annu. Rev. Fluid Mech. 50, 459–482. 10.1146/annurev-fluid-122316-045259.

6. Scallan, J.P., Zawieja, S.D., Castorena-Gonzalez, J.A., and Davis, M.J. (2016). Lymphatic pumping: mechanics, mechanisms and malfunction. J. Physiol. 594, 5749–5768. 10.1113/JP272088.

7. Wigle, J.T., and Oliver, G. (1999). Prox1 Function Is Required for the Development of the Murine Lymphatic System. Cell 98, 769–778. 10.1016/S0092-8674(00)81511-1.

8. Karkkainen, M.J., Haiko, P., Sainio, K., Partanen, J., Taipale, J., Petrova, T.V., Jeltsch, M., Jackson, D.G., Talikka, M., Rauvala, H., et al. (2004). Vascular endothelial growth factor C is required for sprouting of the first lymphatic vessels from embryonic veins. Nat. Immunol. 5, 74–80. 10.1038/ni1013.

9. Yao, L.-C., Baluk, P., Srinivasan, R.S., Oliver, G., and McDonald, D.M. (2012). Plasticity of Button-Like Junctions in the Endothelium of Airway Lymphatics in Development and Inflammation. Am. J. Pathol. 180, 2561–2575. 10.1016/j.ajpath.2012.02.019.

10. Sabine, A., Agalarov, Y., Maby-El Hajjami, H., Jaquet, M., Hägerling, R., Pollmann, C., Bebber, D., Pfenniger, A., Miura, N., Dormond, O., et al. (2012). Mechanotransduction, PROX1, and FOXC2 Cooperate to Control Connexin37 and Calcineurin during Lymphatic-Valve Formation. Dev. Cell 22, 430–445. 10.1016/j.devcel.2011.12.020.

11. Nonomura, K., Lukacs, V., Sweet, D.T., Goddard, L.M., Kanie, A., Whitwam, T., Ranade, S.S., Fujimori, T., Kahn, M.L., and Patapoutian, A. (2018). Mechanically activated ion channel PIEZO1 is required for lymphatic valve formation. Proc. Natl. Acad. Sci. 115, 12817–12822. 10.1073/pnas.1817070115.

12. Wang, Y., Baeyens, N., Corti, F., Tanaka, K., Fang, J.S., Zhang, J., Jin, Y., Coon, B., Hirschi, K.K., Schwartz, M.A., et al. (2016). Syndecan 4 controls lymphatic vasculature remodeling during mouse embryonic development. Development 143, 4441–4451. 10.1242/dev.140129.

13. Cha, B., Ho, Y.-C., Geng, X., Mahamud, Md.R., Chen, L., Kim, Y., Choi, D., Kim, T.H., Randolph, G.J., Cao, X., et al. (2020). YAP and TAZ maintain PROX1 expression in the developing lymphatic and lymphovenous valves in response to VEGF-C signaling. Development. 10.1242/dev.195453.

14. Srinivasan, R.S., Escobedo, N., Yang, Y., Interiano, A., Dillard, M.E., Finkelstein, D., Mukatira, S., Gil, H.J., Nurmi, H., Alitalo, K., et al. (2014). The Prox1–Vegfr3 feedback loop maintains the identity and the number of lymphatic endothelial cell progenitors. Genes Dev. 28, 2175–2187. 10.1101/gad.216226.113.

15. Michalaki, E., Surya, V.N., Fuller, G.G., and Dunn, A.R. (2020). Perpendicular alignment of lymphatic endothelial cells in response to spatial gradients in wall shear stress. Commun. Biol. 3, 57. 10.1038/s42003-019-0732-8.

16. Gong, M.M., Lugo-Cintron, K.M., White, B.R., Kerr, S.C., Harari, P.M., and Beebe, D.J. (2019). Human organotypic lymphatic vessel model elucidates microenvironment-dependent signaling and barrier function. Biomaterials 214, 119225. 10.1016/j.biomaterials.2019.119225.

17. Michalaki, E., Surya, V.N., Rodríguez-Hakim, M., Fuller, G.G., and Dunn, A.R. (2023). Response of lymphatic endothelial cells to combined spatial and temporal variations in fluid flow. FASEB J. 37, e23240. 10.1096/fj.201902205RRRR.

18. Baeyens, N., Nicoli, S., Coon, B.G., Ross, T.D., Van Den Dries, K., Han, J., Lauridsen, H.M., Mejean, C.O., Eichmann, A., Thomas, J.-L., et al. (2015). Vascular remodeling is governed by a VEGFR3-dependent fluid shear stress set point. eLife 4, e04645. 10.7554/eLife.04645.

19. Kuonqui, K.G., Campbell, A.-C., Pollack, B.L., Shin, J., Sarker, A., Brown, S., Park, H.J., Mehrara, B.J., and Kataru, R.P. (2025). Regulation of VEGFR3 signaling in lymphatic endothelial cells. Front. Cell Dev. Biol. 13, 1527971. 10.3389/fcell.2025.1527971.

20. Zhang, H., Rahman, T., Lu, S., Adam, A.P., and Wan, L.Q. (2024). Helical vasculogenesis driven by cell chirality. Sci. Adv. 10, eadj3582. 10.1126/sciadv.adj3582.

21. Ilan, I.S., Yslas, A.R., Peng, Y., Lu, R., and Lee, E. (2023). A 3D Human Lymphatic Vessel-on-Chip Reveals the Roles of Interstitial Flow and VEGF-A/C for Lymphatic Sprouting and Discontinuous Junction Formation. Cell. Mol. Bioeng. 16, 325–339. 10.1007/s12195-023-00780-0.

22. Ayuso, J.M., Gong, M.M., Skala, M.C., Harari, P.M., and Beebe, D.J. (2020). Human Tumor-Lymphatic Microfluidic Model Reveals Differential Conditioning of Lymphatic Vessels by Breast Cancer Cells. Adv. Healthc. Mater. 9, 1900925. 10.1002/adhm.201900925.

23. Lee, E., Chan, S.-L., Lee, Y., Polacheck, W.J., Kwak, S., Wen, A., Nguyen, D.-H.T., Kutys, M.L., Alimperti, S., Kolarzyk, A.M., et al. (2023). A 3D biomimetic model of lymphatics reveals cell–cell junction tightening and lymphedema via a cytokine-induced ROCK2/JAM-A complex. Proc. Natl. Acad. Sci. 120, e2308941120. 10.1073/pnas.2308941120.

24. Cursiefen, C., Chen, L., Borges, L.P., Jackson, D., Cao, J., Radziejewski, C., D’Amore, P.A., Dana, M.R., Wiegand, S.J., and Streilein, J.W. (2004). VEGF-A stimulates lymphangiogenesis and hemangiogenesis in inflammatory neovascularization via macrophage recruitment. J. Clin. Invest. 113, 1040–1050. 10.1172/JCI20465.

25. Cao, R., Eriksson, A., Kubo, H., Alitalo, K., Cao, Y., and Thyberg, J. (2004). Comparative evaluation of FGF-2-, VEGF-A-, and VEGF-C-induced angiogenesis, lymphangiogenesis, vascular fenestrations, and permeability. Circ. Res. 94, 664–670. 10.1161/01.RES.0000118600.91698.BB.

26. Ferrara, N. (1999). Role of vascular endothelial growth factor in the regulation of angiogenesis. Kidney Int. 56, 794–814. 10.1046/j.1523-1755.1999.00610.x.

27. Nakajima, T., Sasaki, K., Yamamori, A., Sakurai, K., Miyata, K., Watanabe, T., and Matsunaga, Y.T. (2020). A simple three-dimensional gut model constructed in a restricted ductal microspace induces intestinal epithelial cell integrity and facilitates absorption assays. Biomater. Sci. 8, 5615– 5627. 10.1039/D0BM00763C.

28. Pauty, J., Usuba, R., Takahashi, H., Suehiro, J., Fujisawa, K., Yano, K., Nishizawa, T., and Matsunaga, Y.T. (2017). A vascular permeability assay using an in vitro human microvessel model mimicking the inflammatory condition. Nanotheranostics 1, 103.

29. Yoshimatsu, Y., Kimuro, S., Pauty, J., Takagaki, K., Nomiyama, S., Inagawa, A., Maeda, K., Podyma-Inoue, K.A., Kajiya, K., Matsunaga, Y.T., et al. (2020). TGF-beta and TNF-alpha cooperatively induce mesenchymal transition of lymphatic endothelial cells via activation of Activin signals. PLOS ONE 15, e0232356. 10.1371/journal.pone.0232356.

30. Price, G.M., Chrobak, K.M., and Tien, J. (2008). Effect of cyclic AMP on barrier function of human lymphatic microvascular tubes. Microvasc. Res. 76, 46–51. 10.1016/j.mvr.2008.02.003.

31. Kim, S., Chung, M., and Jeon, N.L. (2016). Three-dimensional biomimetic model to reconstitute sprouting lymphangiogenesis in vitro. Biomaterials 78, 115–128. 10.1016/j.biomaterials.2015.11.019.

32. Gibot, L., Galbraith, T., Kloos, B., Das, S., Lacroix, D.A., Auger, F.A., and Skobe, M. (2016). Cell-based approach for 3D reconstruction of lymphatic capillaries in vitro reveals distinct functions of HGF and VEGF-C in lymphangiogenesis. Biomaterials 78, 129–139. 10.1016/j.biomaterials.2015.11.027.

33. Osaki, T., Serrano, J.C., and Kamm, R.D. (2018). Cooperative Effects of Vascular Angiogenesis and Lymphangiogenesis. Regen. Eng. Transl. Med. 4, 120–132. 10.1007/s40883-018-0054-2.

34. Cho, Y., Na, K., Jun, Y., Won, J., Yang, J.H., and Chung, S. (2021). Three-Dimensional In Vitro Lymphangiogenesis Model in Tumor Microenvironment. Front. Bioeng. Biotechnol. 9. 10.3389/fbioe.2021.697657.

35. Makinen, T. (2001). Isolated lymphatic endothelial cells transduce growth, survival and migratory signals via the VEGF-C/D receptor VEGFR-3. EMBO J. 20, 4762–4773. 10.1093/emboj/20.17.4762.

36. Breslin, J.W., Yuan, S.Y., and Wu, M.H. (2007). VEGF-C Alters Barrier Function of Cultured Lymphatic Endothelial Cells Through a VEGFR-3-Dependent Mechanism. Lymphat. Res. Biol. 5, 105–114. 10.1089/lrb.2007.1004.

37. Cacheux, J., Nakajima, T., Alcaide, D., Sano, T., Doi, K., Bancaud, A., and Matsunaga, Y.T. (2024). Protocol for fabricating and characterizing microvessel-on-a-chip for human umbilical vein endothelial cells. STAR Protoc. 5, 102950. 10.1016/j.xpro.2024.102950.

38. Wolde-Kidan, A., Herrmann, A., Prause, A., Gradzielski, M., Haag, R., Block, S., and Netz, R.R. (2021). Particle Diffusivity and Free-Energy Profiles in Hydrogels from Time-Resolved Penetration Data. Biophys. J. 120, 463–475. 10.1016/j.bpj.2020.12.020.

39. Cacheux, J., Bancaud, A., Alcaide, D., Suehiro, J.-I., Akimoto, Y., Sakurai, H., and Matsunaga, Y.T. (2023). Endothelial tissue remodeling induced by intraluminal pressure enhances paracellular solute transport. iScience 26, 107141. 10.1016/j.isci.2023.107141.

40. Claesson-Welsh, L., Dejana, E., and McDonald, D.M. (2021). Permeability of the Endothelial Barrier: Identifying and Reconciling Controversies. Trends Mol. Med. 27, 314–331. 10.1016/j.molmed.2020.11.006.

41. Zhang, F., Zarkada, G., Han, J., Li, J., Dubrac, A., Ola, R., Genet, G., Boyé, K., Michon, P., Künzel, S.E., et al. (2018). Lacteal junction zippering protects against diet-induced obesity. Science 361, 599–603. 10.1126/science.aap9331.

42. Zarkada, G., Chen, X., Zhou, X., Lange, M., Zeng, L., Lv, W., Zhang, X., Li, Y., Zhou, W., Liu, K., et al. (2023). Chylomicrons Regulate Lacteal Permeability and Intestinal Lipid Absorption. Circ. Res. 133, 333–349. 10.1161/CIRCRESAHA.123.322607.

43. Yadav, S., and Linstedt, A.D. (2011). Golgi Positioning. Cold Spring Harb. Perspect. Biol. 3, a005322–a005322. 10.1101/cshperspect.a005322.

44. Khuntia, P., and Das, T. (2024). Prediction of Golgi Polarity in Collectively Migrating Epithelial Cells Using Graph Neural Network. Cells Tissues Organs 213, 108–119. 10.1159/000528354.

45. Imoukhuede, P.I., and Popel, A.S. (2011). Quantification and cell-to-cell variation of vascular endothelial growth factor receptors. Exp. Cell Res. 317, 955–965. 10.1016/j.yexcr.2010.12.014.

46. Secker, G.A., and Harvey, N.L. (2021). Regulation of VEGFR Signalling in Lymphatic Vascular Development and Disease: An Update. Int. J. Mol. Sci. 22, 7760. 10.3390/ijms22147760.

47. Nilsson, I., Bahram, F., Li, X., Gualandi, L., Koch, S., Jarvius, M., Söderberg, O., Anisimov, A., Kholová, I., Pytowski, B., et al. (2010). VEGF receptor 2/-3 heterodimers detected in situ by proximity ligation on angiogenic sprouts. EMBO J. 29, 1377–1388. 10.1038/emboj.2010.30.

48. Joukov, V., Kumar, V., Sorsa, T., Arighi, E., Weich, H., Saksela, O., and Alitalo, K. (1998). A Recombinant Mutant Vascular Endothelial Growth Factor-C that Has Lost Vascular Endothelial Growth Factor Receptor-2 Binding, Activation, and Vascular Permeability Activities. J. Biol. Chem. 273, 6599–6602. 10.1074/jbc.273.12.6599.

49. Millán, J., Cain, R.J., Reglero-Real, N., Bigarella, C., Marcos-Ramiro, B., Fernández-Martín, L., Correas, I., and Ridley, A.J. (2010). Adherens junctions connect stress fibres between adjacent endothelial cells. BMC Biol. 8, 11. 10.1186/1741-7007-8-11.

50. Heinolainen, K., Karaman, S., D’Amico, G., Tammela, T., Sormunen, R., Eklund, L., Alitalo, K., and Zarkada, G. (2017). VEGFR3 Modulates Vascular Permeability by Controlling VEGF/VEGFR2 Signaling. Circ. Res. 120, 1414–1425. 10.1161/CIRCRESAHA.116.310477.

51. Deng, Y., Zhang, X., and Simons, M. (2015). Molecular Controls of Lymphatic VEGFR3 Signaling. Arterioscler. Thromb. Vasc. Biol. 35, 421–429. 10.1161/ATVBAHA.114.304881.

52. Dixelius, J., Mäkinen, T., Wirzenius, M., Karkkainen, M.J., Wernstedt, C., Alitalo, K., and Claesson-Welsh, L. (2003). Ligand-induced Vascular Endothelial Growth Factor Receptor-3 (VEGFR-3) Heterodimerization with VEGFR-2 in Primary Lymphatic Endothelial Cells Regulates Tyrosine Phosphorylation Sites. J. Biol. Chem. 278, 40973–40979. 10.1074/jbc.M304499200.

53. Blake, R.A., Broome, M.A., Liu, X., Wu, J., Gishizky, M., Sun, L., and Courtneidge, S.A. (2000). SU6656, a Selective Src Family Kinase Inhibitor, Used To Probe Growth Factor Signaling. Mol. Cell. Biol. 20, 9018–9027. 10.1128/MCB.20.23.9018-9027.2000.

54. Cheng, M., Wu, J., Li, Y., Nie, Y., and Chen, H. (2008). Activation of MAPK participates in low shear stress-induced IL-8 gene expression in endothelial cells. Clin. Biomech. 23, S96–S103. 10.1016/j.clinbiomech.2008.06.003.

55. Leelahavanichkul, K., Amornphimoltham, P., Molinolo, A.A., Basile, J.R., Koontongkaew, S., and Gutkind, J.S. (2014). A role for p38 MAPK in head and neck cancer cell growth and tumor-induced angiogenesis and lymphangiogenesis. Mol. Oncol. 8, 105–118. 10.1016/j.molonc.2013.10.003.

56. Breslin, J.W. (2011). ROCK and cAMP Promote Lymphatic Endothelial Cell Barrier Integrity and Modulate Histamine and Thrombin-Induced Barrier Dysfunction. Lymphat. Res. Biol. 9, 3–11. 10.1089/lrb.2010.0016.

57. Lång, E., Lång, A., Blicher, P., Rognes, T., Dommersnes, P.G., and Bøe, S.O. (2024). Topology-guided polar ordering of collective cell migration. Sci. Adv. 10, eadk4825. 10.1126/sciadv.adk4825.

58. Lin, S.-Z., Ye, S., Xu, G.-K., Li, B., and Feng, X.-Q. (2018). Dynamic Migration Modes of Collective Cells. Biophys. J. 115, 1826–1835. 10.1016/j.bpj.2018.09.010.

59. Lo, C.M., Wang, H.B., Dembo, M., and Wang, Y.L. (2000). Cell movement is guided by the rigidity of the substrate. Biophys. J. 79, 144–152. 10.1016/S0006-3495(00)76279-5.

60. Saez, A., Ghibaudo, M., Buguin, A., Silberzan, P., and Ladoux, B. (2007). Rigidity-driven growth and migration of epithelial cells on microstructured anisotropic substrates. Proc. Natl. Acad. Sci. 104, 8281–8286. 10.1073/pnas.0702259104.

61. Skillin, N.P., Kirkpatrick, B.E., Herbert, K.M., Nelson, B.R., Hach, G.K., Günay, K.A., Khan, R.M., DelRio, F.W., White, T.J., and Anseth, K.S. (2024). Stiffness anisotropy coordinates supracellular contractility driving long-range myotube-ECM alignment. Sci. Adv. 10, eadn0235. 10.1126/sciadv.adn0235.

62. Lee, H.-H., Tien, S.-C., Jou, T.-S., Chang, Y.-C., Jhong, J.-G., and Chang, Z.-F. (2010). Src-dependent phosphorylation of ROCK participates in regulation of focal adhesion dynamics. J. Cell Sci. 123, 3368–3377. 10.1242/jcs.071555.

63. Ueda, M., Terai, Y., Yamashita, Y., Kumagai, K., Ueki, K., Yamaguchi, H., Akise, D., Hung, Y., and Ueki, M. (2002). Correlation between vascular endothelial growth factor-C expression and invasion phenotype in cervical carcinomas. Int. J. Cancer 98, 335–343. 10.1002/ijc.10193.

64. Hall, E., Mendiola, K., Lightsey, N.K., and Hanjaya-Putra, D. (2024). Mimicking blood and lymphatic vasculatures using microfluidic systems. Biomicrofluidics 18, 031502. 10.1063/5.0175154.

65. Zhong, J., Shelton, E.L., Kirabo, A., and Kon, V. (2023). Mechanisms of Lymphatic Endothelial Cell Junction Transformations. Circ. Res. 133, 350–352. 10.1161/CIRCRESAHA.123.323210.

66. Cacheux, J., Ordonez-Miranda, J., Bancaud, A., Jalabert, L., Alcaide, D., Nomura, M., and Matsunaga, Y.T. (2023). Asymmetry of tensile versus compressive elasticity and permeability contributes to the regulation of exchanges in collagen gels. Sci. Adv. 9, eadf9775. 10.1126/sciadv.adf9775.

67. Sato, H., Katano, M., Takigawa, T., and Masuda, T. (2001). Estimation for the elasticity of vascular endothelial cells on the basis of atomic force microscopy and Young’s modulus of gelatin gels. Polym. Bull. 47, 375–381. 10.1007/s289-001-8195-z.

68. Ohashi, T., Ishii, Y., Ishikawa, Y., Matsumoto, T., and Sato, M. (2002). Experimental and numerical analyses of local mechanical properties measured by atomic force microscopy for sheared endothelial cells. Biomed. Mater. Eng. 12, 319–327.

69. Bastounis, E.E., Yeh, Y.-T., and Theriot, J.A. (2019). Subendothelial stiffness alters endothelial cell traction force generation while exerting a minimal effect on the transcriptome. Sci. Rep. 9, 18209. 10.1038/s41598-019-54336-2.

70. Schindelin, J., Arganda-Carreras, I., Frise, E., Kaynig, V., Longair, M., Pietzsch, T., Preibisch, S., Rueden, C., Saalfeld, S., Schmid, B., et al. (2012). Fiji: an open-source platform for biological-image analysis. Nat. Methods *9*, 676–682. 10.1038/nmeth.2019.

71. Arganda-Carreras, I., Kaynig, V., Rueden, C., Eliceiri, K.W., Schindelin, J., Cardona, A., and Sebastian Seung, H. (2017). Trainable Weka Segmentation: a machine learning tool for microscopy pixel classification. Bioinformatics 33, 2424–2426. 10.1093/bioinformatics/btx180.

72. Schmidt, U., Weigert, M., Broaddus, C., and Myers, G. (2018). Cell Detection with Star-Convex Polygons. In Medical Image Computing and Computer Assisted Intervention – MICCAI 2018 Lecture Notes in Computer Science., A. F. Frangi, J. A. Schnabel, C. Davatzikos, C. Alberola-López, and G. Fichtinger, eds. (Springer International Publishing), pp. 265–273. 10.1007/978-3-030-00934-2_30.

73. Ershov, D., Phan, M.-S., Pylvänäinen, J.W., Rigaud, S.U., Le Blanc, L., Charles-Orszag, A., Conway, J.R.W., Laine, R.F., Roy, N.H., Bonazzi, D., et al. (2022). TrackMate 7: integrating state-of-the-art segmentation algorithms into tracking pipelines. Nat. Methods 19, 829–832. 10.1038/s41592-022-01507-1.

74. Tinevez, J.-Y., Perry, N., Schindelin, J., Hoopes, G.M., Reynolds, G.D., Laplantine, E., Bednarek, S.Y., Shorte, S.L., and Eliceiri, K.W. (2017). TrackMate: an open and extensible platform for single-particle tracking. Methods 115, 80–90. 10.1016/j.ymeth.2016.09.016.

75. Wortel, I.M.N., Liu, A.Y., Dannenberg, K., Berry, J.C., Miller, M.J., and Textor, J. (2021). CelltrackR: An R package for fast and flexible analysis of immune cell migration data. ImmunoInformatics 1–2, 100003. 10.1016/j.immuno.2021.100003.

76. Waskom, M. (2021). seaborn: statistical data visualization. J. Open Source Softw. 6, 3021. 10.21105/joss.03021.

